# ANGPTL3 regulates the peroxisomal translocation of SmarcAL1 in response to cell growth states

**DOI:** 10.1101/2024.06.03.597253

**Authors:** Taylor Hanta Nagai, Taiji Mizoguchi, Yanyan Wang, Amy Deik, Kevin Bullock, Clary B. Clish, Yu-Xin Xu

## Abstract

Angiopoietin-like 3 (ANGPTL3) is a key regulator of lipoprotein metabolism, known for its potent inhibition on intravascular lipoprotein and endothelial lipase activities. Recent studies have shed light on the cellular functions of ANGPTL3. However, the precise mechanism underlying its regulation of cellular lipid metabolism remains elusive. We recently reported that ANGPTL3 interacts with the chromatin regulator SMARCAL1, which plays a pivotal role in maintaining cellular lipid homeostasis. Here, through a combination of in vitro and in vivo functional analyses, we provide evidence that ANGPTL3 indeed influences cellular lipid metabolism. Increased expression of Angptl3 prompted the formation of lipid droplets (LDs) in response to slow growth conditions. Notably, under the conditions, Angptl3 accumulated within cytoplasmic peroxisomes, where it interacts with SmarcAL1, which translocated from nucleus as observed previously. This translocation induced changes in gene expression favoring triglyceride (TG) accumulation. Indeed, *ANGPTL3* gene knockout (KO) in human cells increased the expression of key lipid genes, which could be linked to elevated nuclear localization of SMARCAL1, whereas the expression of these genes decreased in *SMARCAL1* KO cells. Consistent with these findings, the injection of Angptl3 protein to mice led to hepatic fat accumulation derived from circulating blood, a phenotype likely indicative of its long-term effect on blood TG, linked to SmarcAL1 activities. Thus, our results suggest that the Angptl3-SmarcAL1 pathway may confer the capacity for TG storage in cells in response to varying growth states, which may have broad implications for this pathway in regulating energy storage and trafficking.

## Introduction

Angiopoietin-like 3 (ANGPTL3) belongs to the family of angiopoietin-like proteins (ANGPTLs). The first evidence that Angptl3 is involved in lipid metabolism came from a genetic study in mice ^1^. Obese (*ob/ob*) mice normally display high levels of plasma insulin, glucose, and lipids. One mutant strain (*KK/San*) derived from the *ob/ob* mice exhibited an opposite phenotype - low plasma lipids and particularly very low triglyceride (TG); a null mutation in the *Angptl3* coding region was found to be responsible for this phenotype.

Human genetic studies demonstrated that ANGPTL3 is a key lipoprotein regulator ^2,3^. *ANGPTL3* null mutations cause familial combined hypolipidemia ^3^. Individuals deficient in *ANGPTL3* (due to compound heterozygosity for two null mutations) showed extremely low plasma levels of not only TG, but also low-density lipoprotein cholesterol (LDL-C) and high-density lipoprotein cholesterol (HDL-C). Population-based studies show that *ANGPTL3* deficiency protects against the development of coronary artery disease (CAD) ^4^, providing strong evidence that ANGPTL3 could be a therapeutic target for treatment of dyslipidemia.

ANGPTL3 is a secreted protein that is exclusively expressed in the livers ^5^. It has an N-terminal coiled-coil domain and a C-terminal fibrinogen domain. The intact protein can be cleaved at the linker region between the two domains ^6,7^. Most of the studies on ANGPTL3 have focused on its intravascular activities for its inhibition of lipoprotein lipase (LPL) ^7,8^, and endothelial lipase ^9^. For instance, the inhibition of LPL was shown to be crucial in the clearance of TG from chylomicrons and very low-density lipoprotein (VLDL) particles or TG-rich lipoproteins (TGLs) ^10,11^. The inhibition of LPL by Angptl3 likely explains the low TG observed in humans and mice carrying *Angptl3* null mutations. Therefore, this inhibition is thought to be the major mechanism of ANGPTL3 in regulating TG metabolism. It should be noted that the current mechanism based on LPL inhibition can explain some, but not all, of the observed lipid phenotypes ^12,13^.

Therapeutic inhibition of ANGPTL3 has emerged as a promising strategy for treating dyslipidemia and reducing the risk of CAD and metabolic disorders ^14–16^. For instance, inhibition of ANGPTL3 expression with anti-sense oligonucleotides (ASOs) has been shown to significantly reduce levels of TG and cholesterols in both mice and humans, leading to beneficial cardiometabolic effects ^16^. Clinical trial employing this approach has effectively lowered the non-HDL-C levels in statin-treated patients with elevated cholesterol. However, this trial was prematurely terminated due to hepatic fat accumulation resulting from ASO-mediated silencing of ANGPTL3 expression ^17^. Prior to this clinical trial, we observed TG accumulation in human cells lacking ANGPTL3, and silencing ANGPTL3 expression with siRNA decreased apolipoprotein (Apo) B (or VLDL) secretion ^18^. Recent studies further support these observations, underscoring the critical roles of proper ANGPTL3 expression in lipid metabolism (see more in discussion) ^19,20^. In particular, ANGPTL3 silencing with siRNAs induced fat accumulation, potentially linked to cellular energy metabolism ^20^. While initial evidence predominantly highlights the cellular functions of ANGPTL3, there remains a pressing need to investigate how ANGPTL3 precisely regulates cellular lipid metabolism.

In this study, we provided evidence that Angptl3 modulates cellular lipid metabolism. We found that, under slow growth conditions, the increased expression of Angptl3 led to a significant rise in lipid droplet (LD) formation that is directly linked to cellular accumulation of TGs and fatty acids (FAs). Notably, we observed that under these slow growth conditions, cytoplasmic Angptl3 was redistributed, becoming enriched on peroxisomes where it co-localized with the nuclear chromatin regulator SmarcAL1. This translocation of SmarcAL1 from the nucleus, as previously reported ^21^, had a profound impact on the regulation of gene expression in response to growth conditions. Specifically, it resulted in the down-regulation of a considerable number of genes essential for lipid metabolism, thereby favoring the accumulation of TGs and FAs. Our in vivo analysis further supports these findings. Injection of Angptl3 protein in mice effectively lowered blood TG levels and increased fat accumulation in the livers consistent with SmarcAL1 activities. These results collectively indicate that Angptl3-regulated SmarcAL1 activity is a crucial factor in regulating cellular lipid metabolism in response to varying cell growth states, suggesting that this pathway may hold broad significance in mediating systemic energy storage.

## Results

### Angptl3 modulates cellular lipid metabolism

We recently reported the interaction of Angptl3 with the chromatin regulator SmarcAL1 from our proteomics analysis based on rat liver cell lines, McArdle-RH7777 (McA), that stably expressing a Fc tag or Fc-tagged Angptl3 ^21^. The interaction with SmarcAL1 suggested that Angptl3 might be involved in cellular lipid metabolism. To gain direct insights into the significance of the interaction, we conducted more functional analyses using these cell lines. Under normal growth condition with 10% FBS, we observed a marginal increase in the number of LDs in Fc-Angptl3 cells (Fig. 1A, top). Intriguingly, when the cells were cultured in low serum media with 2% FBS, a substantial surge in LDs was particularly evident in Fc-Angptl3 cells, exhibiting an ∼50-fold increase (Fig. 1A, bottom). We reasoned that the increased LD formation might be related to the slowed cell growth. To investigate this hypothesis, we subjected the cells to lower temperatures (30 °C) for further slower growth. Remarkably, we observed the emergence of massive LDs exclusively in Fc- Angptl3 cells, not only in low serum media but also in cells cultured with 10% FBS (Fig. 1B).

**Fig. 1.**
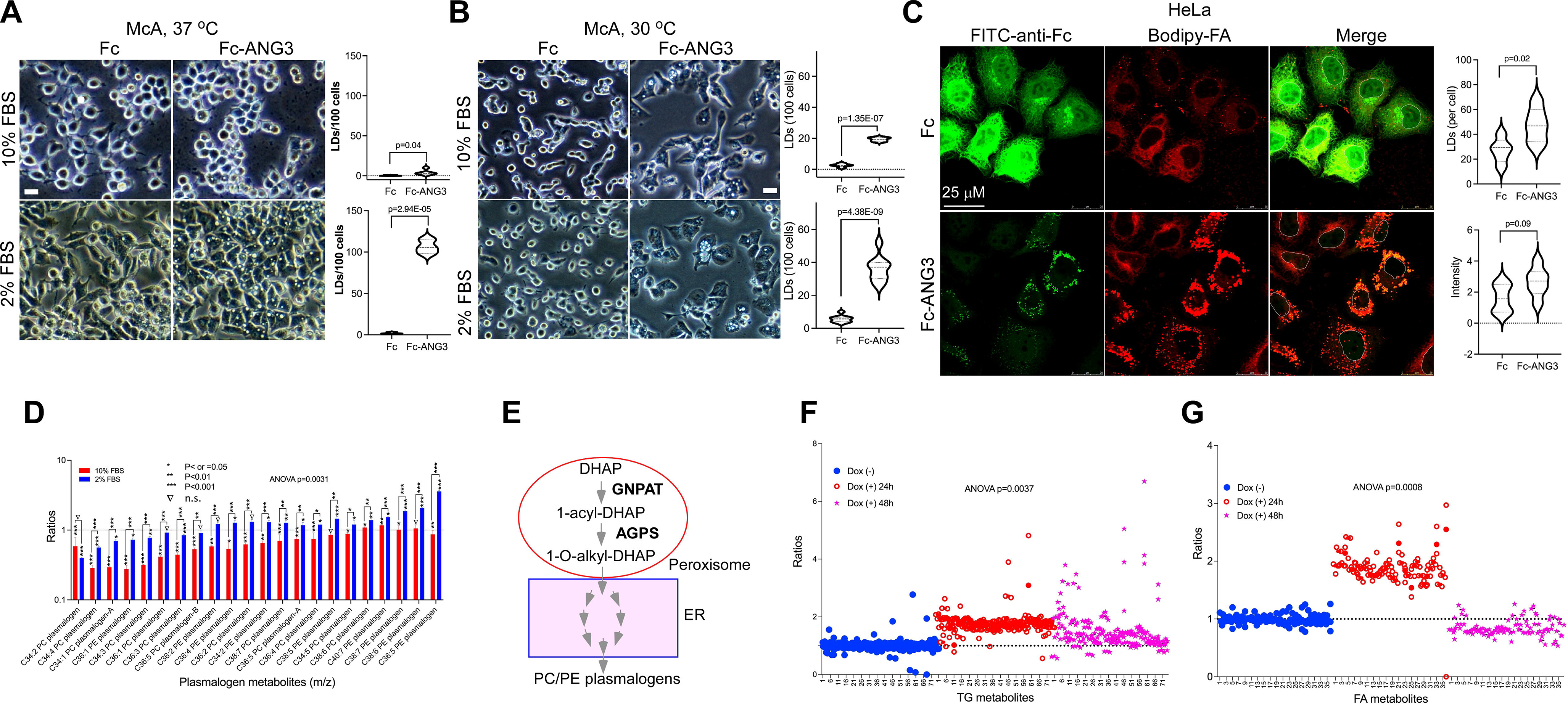
**Increased Angptl3 expression affects cellular lipid metabolism**. **A**. and **B**. Angptl3 expression induces LD formation. Fc and Fc-Angptl3 cells were grown in DMEM media with 10% FBS or OPTI-MEM media with 2% FBS at 37 °C (A) or 30 °C (B) for three days. Phase images were obtained from light microscopy. Quantification was from three independent experiments (right). **C**. Transient Angptl3 expression is sufficient to induce LD formation. HeLa cells transfected with Fc or Fc-Angptl3 plasmids were labeled with BODIPY 558/568 C12 and stained with FITC-labeled anti-Fc antibody. Images were obtained from confocal microscopy. **D**. Comparative analysis of plasmalogen metabolite changes. Total lipid and polar lipid extracts from Fc and Fc-Angptl3 cells as in A were analyzed with mass spectrometry (see sFig. 2 and sTable 1 for details). Ratios were calculated by comparing metabolites from Fc-Angptl3 cells with those from Fc cells. p values were calculated by comparing Fc-Angptl3 cells with Fc cells under each or both growth conditions. **E**. Diagram of plasmalogen biosynthesis pathway. **F**. Inducible expression of Angptl3 transiently elevated TG and FA levels. Cell lines carrying inducible Fc-Angptl3 constructs were treated with or without doxycycline (Dox) for 24 and 48 hrs. Total lipid and polar lipid extracts were analyzed with mass spectrometry. Ratios were calculated by comparing metabolites with the averages of those from three replicates without Dox treatment (see sFig. 3 and sTable 2 for details). P values calculated from two-sided unpaired t-test or two-way ANOVA as indicated. Bars, 25 μM.

The results suggest that elevated Angptl3 expression has an impact on the distribution of cellular lipids. To validate this lipid phenotype across different cell types, we conducted transient expression experiments involving Fc and Fc-Angptl3 in HeLa cells. The results reveal that the presence of Fc-Angptl3, as opposed to Fc alone, induces a noticeable alteration in bodipy-labeled LD formation (Fig. 1C, bottom). Comparable results were observed in McA cells, either tagged with Fc or GFP tags (sFig. 1A and 1B). The expression of the N-terminal domain of Angptl3 alone proves sufficient to elicit a similar effect, albeit at a lower expression efficiency (sFig. 1B and 1C, bottom).

To examine the cellular lipid metabolite changes, we performed a metabolomics analysis of the stable cell lines under the low serum condition. The results reveal no consistent changes in neutral lipids and FAs (sFig. 2A-2D and sTable 1). Notably, an increase in lipids with shorter carbon chains, including monoglycerides (MGs), diglycerides (DGs), and some TGs, was observed, along with a decrease in FAs and TGs with longer carbon chains specifically in the Fc-Angptl3 cells with 2% FBS (sFig. 2A-2C). Despite these alterations, it appears unlikely that these changes alone would account for the substantial LD formation observed in these cells.

**Fig. 2.**
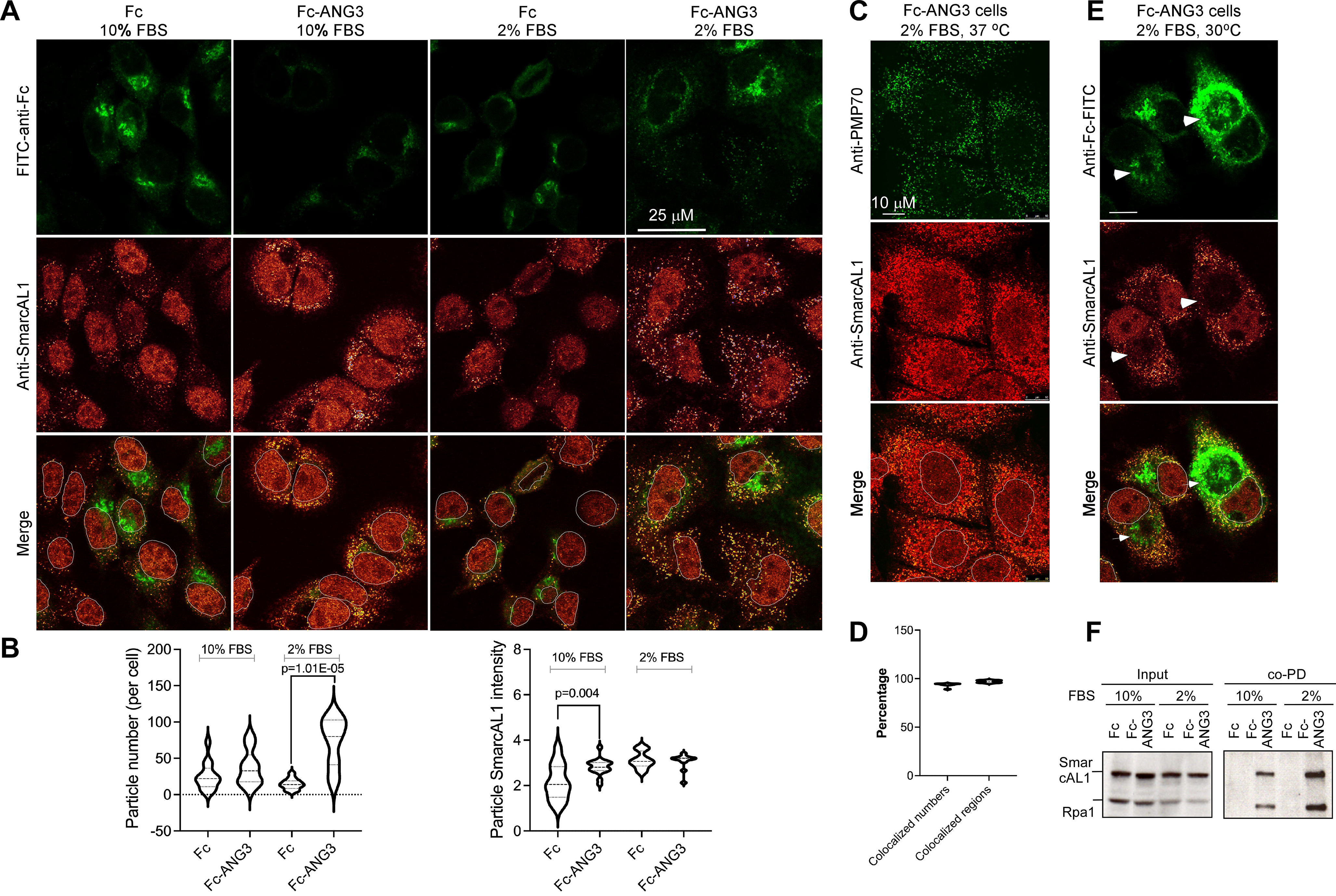
**Angptl3 regulates peroxisomal translocation of SmarcAL1**. **A**. Cytoplasmic enrichment of SmarcAL1. Fc and Fc-Angptl3 McA cells grown with 10% or 2% FBS media at 37 °C for two days were fixed and hybridized with FITC-labeled anti-Fc (green), anti- SmarcAL1 (red) plus fluorescence-labeled secondary antibodies. Images were acquired from confocal microscopy **B**. Quantification of cytoplasmic SmarcAL1 localization. Quantification was carried out from three independent experiments using CellProfiler. P values were calculated with two-sided unpaired t-test. **C**. Peroxisomal localization of SmarcAL1. Representative images were obtained as in A, but with Fc-Angptl3 McA cells grown with 2% FBS media at 37 °C for two days hybridized with anti-PMP70 (peroxisome marker, green) and -SmarcAL1 (red) plus fluorescence-labeled secondary antibodies. **D**. Quantification of colocalized particle numbers and regions as in C. **E**. Enrichment of Fc- Angptl3 protein at the periphery of LDs. Representative images were obtained as in A, but from Fc-Angptl3 McA cells with 2% FBS media at 30 °C hybridized with FITC-labeled anti- Fc (green) and anti-SmarcAL1 (red), and fluorescence-labeled secondary antibodies. Arrows indicate the enrichment of Fc-Angptl3 protein at the periphery of large LDs (see sFig. 4 for more details). **F**. Affinity PD analysis of Fc and Fc-Angptl3 McA cells grown in the media with 10% or 2% FBS as in A. The PD products were analyzed with Western blotting probed with anti-SmarcAL1 and -Rpa1 antibodies.

Unexpectedly, serum starvation consistently induced elevated levels of phosphatidylcholine (PC) and phosphatidylethanolamine (PE) plasmalogens (Fig. 1D), along with numerous other phospholipids (sFig. 2E-2G and sTable 1). As reported previously ^22^, increased surface PC or a PC/PE mixture can stabilize LDs by preventing their coalescence. Consequently, the significant upsurge in LD formation may be attributed to the elevated levels of PC/PE plasmalogens in the cells under lower serum conditions. Considering the early biogenesis of plasmalogens at peroxisomes (Fig. 1E), the extensive LD formation under reduced serum conditions may be linked to peroxisome activities (see below). In line with this, the expression of the key peroxisome gene AGPS (fold change,1.23; FDR p value<0.05) was specifically up-regulated under low serum condition ^23^.

The above results provided some direct evidence that Angptl3 is linked to cellular lipid metabolism. However, we realized that stable cells might have undergone many adaptive changes to counteract any adverse effects on growth resulting from Fc-Angptl3 expression. Consequently, the observed cellular metabolite changes might not be the immediate consequences of Angptl3 activities. To address this, we established HepG2 cell lines with inducible expression of Fc or Fc-Angptl3 (sFig. 3). The addition of doxycycline effectively induced the expression of Fc-Angptl3 (due to leaking expression, the Fc expression was not increased upon induction) (sFig. 3A). Transient expression of Fc-Angptl3 robustly elevated the levels of TGs, FAs (see Fig. 1F and 1G and sTable 2), plasmalogens, and various other lipid metabolites (sFig. 3B-3E and sTable 2). Notably, these increases were not observed in the Fc cells (data not shown). The observed metabolite changes were likely triggered by the interaction of Angptl3 with SmarcAL1 (sFig. 3F), a phenotype confirmed in this experimental system, as previously reported ^21^.

**Fig. 3.**
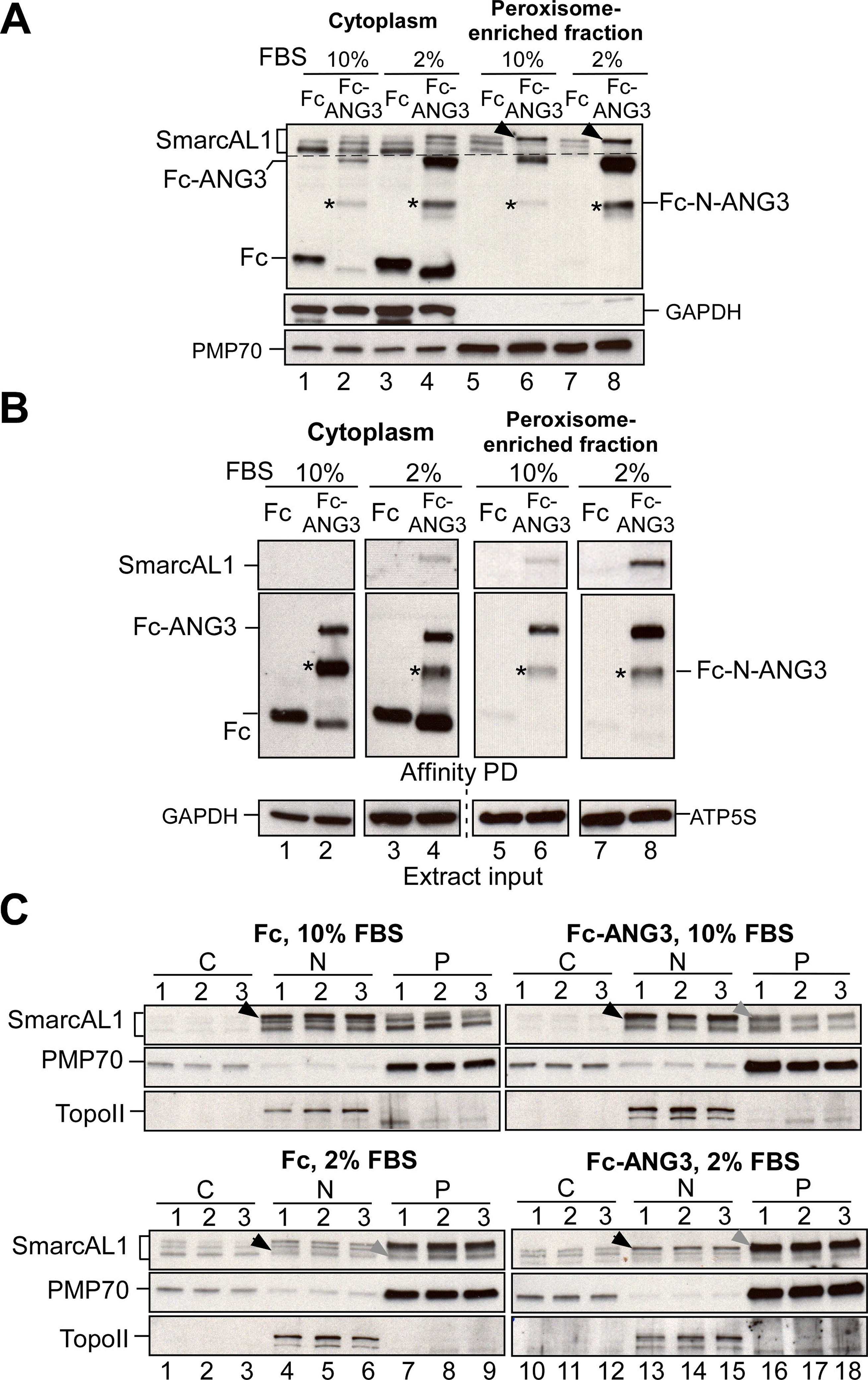
**Biochemical validation of Angptl3-SmarcAL1 interaction at peroxisomes and SmarcAL1’s peroxisomal translocation. A**. SmarcAL1 proteins landed on peroxisomes are highly modified. Soluble cytoplasm and peroxisome-enriched fraction from Fc and Fc- Angptl3 cells grown in media with 10% or 2% FBS were analyzed with Western blotting (with better separation for SmarcAL1) and detected with anti-SmarcAL1, -PMP70 (peroxisome marker), and -GAPDH (cytoplasm marker) antibodies. Heavily modified SmarcAL1 associated with peroxisomes are indicated with arrows. **B**. Serum starvation enhances Angptl3-SmarcAL1 interaction on peroxisomes. Soluble cytoplasm and peroxisome-enriched fractions from Fc and Fc-Angptl3 McA cells with 10% or 2% FBS (as in A and B) was pulled down with protein-A beads and precipitates were detected with anti-SmarcAL1 (top), -Fc (middle) or -GAPDH or -ATP5s (bottom) antibodies. **C.** Serum starvation enhances the translocation of SmarcAL1 to peroxisomes. Soluble cytoplasm (C), nuclear (N) and peroxisome-enriched (P) fractions from Fc (left) or Fc-Angptl3 (right) McA cells with 10% (top) or 2% (bottom) FBS were analyzed with Western blotting and detected with anti-SmarcAL1, -PMP70, and -topoisomerase IIα (TopoII, nuclear marker) antibodies. *, N-terminal Fc-Angptl3.

All these functional analyses collectively suggest that the modulation of Angptl3 expression induces cellular lipid phenotypes resulting from quantitative changes in lipid metabolites. This finding indicates that Angptl3 plays a regulatory role in cellular lipid metabolism.

### Angptl3 regulates the translocation of SmarcAL1 to cytoplasmic peroxisomes

We recently reported the interaction between Angptl3 and SmarcAL1, providing a comprehensive analysis that establishes SmarcAL1’s pivotal role in cellular lipid homeostasis ^21^. The observed lipid phenotypes induced by Angptl3 led us to investigate potential connections with SmarcAL1 activities. Specifically, we sought to explore the distribution of SmarcAL1 in response to increased Angptl3 expression. In our initial investigation, we conducted confocal immunofluorescence assays utilizing established cell lines, as in Figure 1A. The results (Fig. 2A and 2B) revealed that in Fc cells with 10% FBS, a stronger SmarcAL1 signal was observed in the nuclei, accompanied by a weaker signal in sparse dot-like particles likely representing SmarcAL1-enriched areas in the cytoplasm (Fig. 2A, left). In contrast, the Fc-Angptl3 cells under the same conditions exhibited a significant increase in the number of dot-like particles in the cytoplasm (∼2.9- fold increase). Notably, under conditions with 2% FBS (Fig. 2A, right), the Fc-Angptl3 cells displayed a drastic (∼7.7-fold) increase in cytoplasmic dot-like particles. While not universal, a portion of Fc-Angptl3 signals also appeared in dot-like particles, particularly under serum starvation, and these particles showed co-localization with SmarcAL1 (Fig. 2A, right) (see below).

In our previous observations, we noted the translocation of SmarcAL1 from nuclear to cytoplasmic peroxisomes ^21^. The dot-like particles in the cytoplasm resembled those associated with peroxisomes. To validate this observation, we conducted a confocal immunofluorescence analysis using anti-PMP70 (peroxisome marker) ^24^ and anti- SmarcAL1 antibodies in the dot-like particle-enriched Fc-Angptl3 cells cultured with 2% FBS (see below). The analysis confirmed the co-localization of SmarcAL1 within the dot-like particles with PMP70 as in Fig. 1B (Fig. 2C and 2D).

We further examined the cell lines grown at 30°C (sFig. 4). The results revealed a significant increase in the number of dot-like particles containing SmarcAL1, not only in the Fc-Angptl3 cells with low serum but also in cells with 10% FBS, indicating cytoplasmic enrichment of SmarcAL1 under this low-temperature growth condition.

**Fig. 4.**
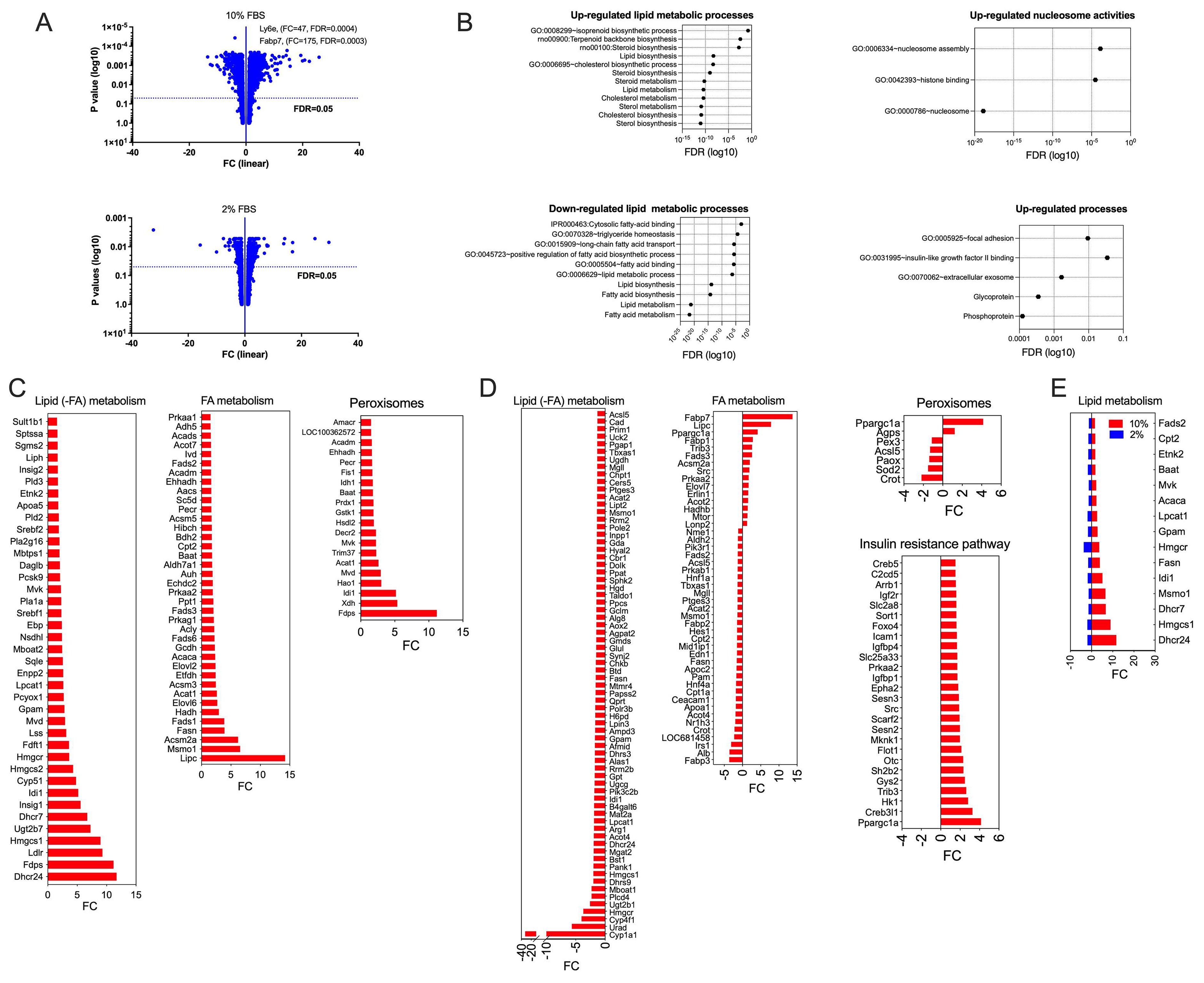
**Gene expression analysis for changes induced by slow growth state**. Total RNA from Fc and Fc-Angptl3 McA cells under normal growth (10% FBS) or serum starvation (2% FBS) was used in rat GeneChip microarray assays. **A**. Volcano plots for differentially expressed genes from the cells with 10% (top) and 2% FBS (bottom). **B**. Gene ontology (GO) enrichment analysis from the cells with 10% (top) and 2% FBS (bottom). **C** and **D**. Differentially expressed genes related to lipid and FA metabolism from the cells with 10% (C) and 2% FBS (D). **E**. The differentially expressed common gene list with up-regulated genes from the cells with 10% FBS and down-regulated in the cells with 2% FBS.

Under this condition, there are multiple phases of Fc-Angptl3 cells (sFig. 4B). In certain cells, Angptl3 is colocalized with SmarcAL1 in the majority of the particles (sFig. 4B, bottom). The most pronounced change observed was the accumulation of a substantial amount of Fc-Angptl3 protein at the periphery of these large LDs in some Fc-Angptl3 cells (Fig. 2D and sFig. 4B, top). While many changes prior to this accumulation might occur, it is essential to highlight that, under the conditions of Angptl3 overexpression and slow cell growth, we observed an increased interaction between Angptl3 and SmarcAL1, along with enhanced SmarcAL1 shuttling to cytoplasmic peroxisomes (Fig. 2F), potentially triggering LD formation.

To assess the dynamic changes of SmarcAL1 in the cytoplasm under the conditions tested, we conducted a time-course assay (sFig. 5). Initially, cells were cultured in normal media, and the time-course initiation occurred following the switch to low serum media. We observed a gradual enrichment of SmarcAL1 at peroxisomes, reaching its peak around 24 hrs and exhibiting a slight decrease around 48 hrs. Interestingly, this enrichment coincided with the appearance of PMP70 during the time course (sFig. 5B and 5C). Importantly, it is worth noting that SmarcAL1 is not essential for peroxisome formation, as evident from the absence of significant changes in peroxisomes in SmarcAL1 KO cells ^21^.

**Fig. 5.**
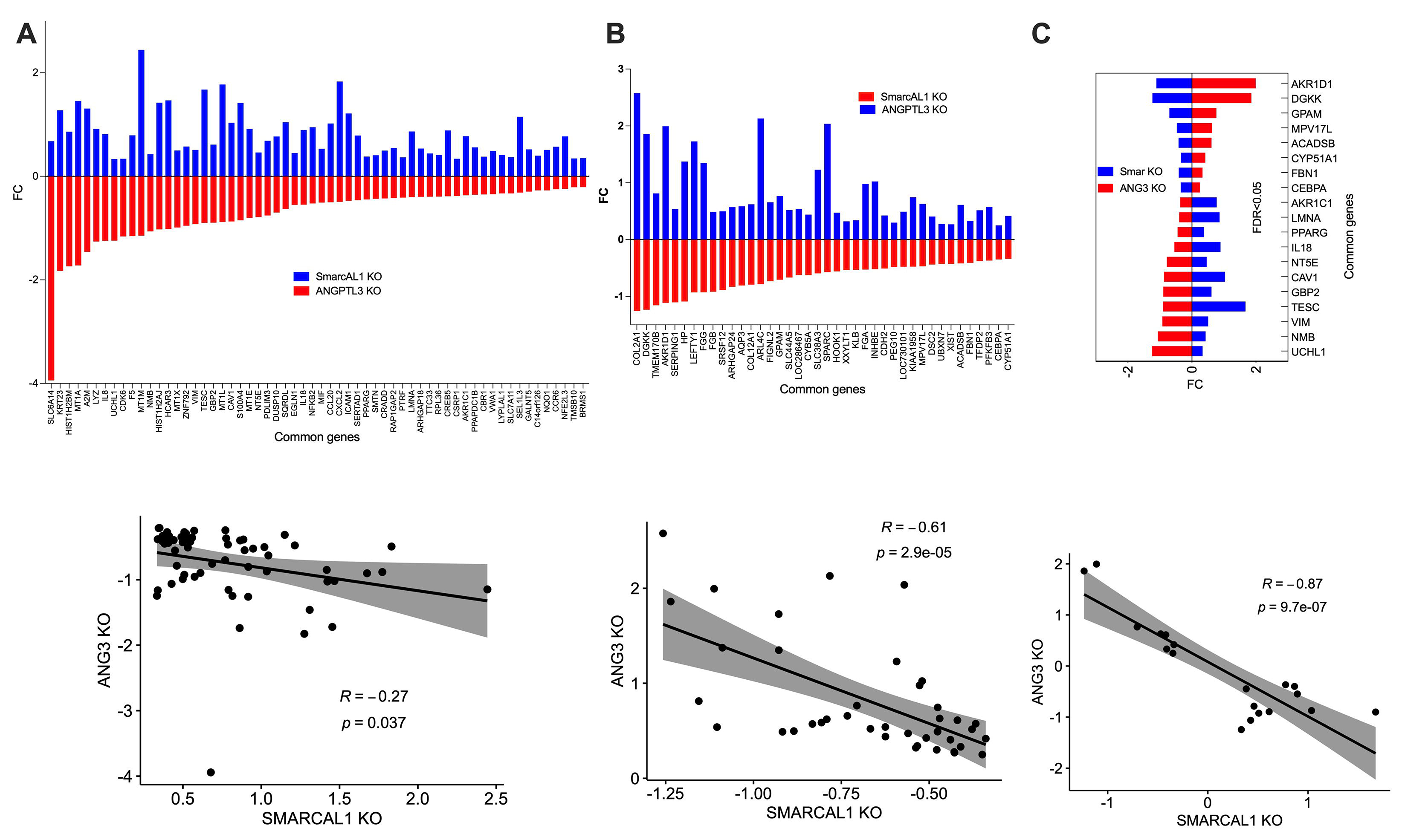
**Comparative analysis of gene expression profiles related to SMARCAL1 activities in SMARCAL1 KO and ANGPTL3 KO Huh7 cells. A**. Expression profiles of genes up-regulated in SMARCAL1 KO and down-regulated in ANGPTL3 KO Huh7 cells. **B**. Expression profiles of genes up-regulated in ANGPTL3 KO and down-regulated in SMARCAL1 KO Huh7 cells. **C**. Shared genes that were up-regulated in ANGPTL3 KO and down-regulated in both SMARCAL1 KO and ANGPTL3 KO Huh7 cells, as well as vice versa. At the bottom of each panel, the correlation results, including regression coefficients and p values from regression analyses for each dataset, are provided.

Subsequently, we employed biochemical analyses to reaffirm SmarcAL1 shuttling between nucleus and cytoplasm based on the cell models under the tested conditions. Extracts were prepared from nuclei (N), soluble cytoplasm (C), and peroxisome-enriched (P) fractions from both Fc and Fc-Angptl3 cells cultured in media with either 10% or 2% FBS (Fig. 3). The extracts were validated using antibodies against markers specific to each cell fraction, as indicated. To explore potential modifications of SmarcAL1 in each fraction, the extracts were initially assessed for changes in SmarcAL1 mobility through Western blot analysis. A notable distinction emerged, revealing a higher abundance of cytoplasmic SmarcAL1 proteins with slower mobility in Fc-Angptl3 cells compared to control cells. Particularly under lower serum conditions, the majority of peroxisomal SmarcAL1 proteins in Fc-Angptl3 cells exhibited slower mobility (Fig. 3A, as indicated with arrows). These modification changes in SmarcAL1 were found to be correlated with Angptl3 levels.

To ascertain the physical interaction between Angptl3 and SmarcAL1 at peroxisomes, we used the soluble cytoplasm and cytoplasmic peroxisome-enriched fraction for affinity pull-down (PD). The results (Fig. 3B) revealed a limited interaction between Angptl3 and SmarcAL1 in the peroxisome fraction under normal growth conditions (lane 6). However, this interaction was significantly enhanced under serum starvation (lane 8). These biochemical analyses confirmed the physical interaction at peroxisomes and aligned closely with the aforementioned immunofluorescence and biochemical analyses (Fig. 2 and 3A). This consistency underscores that SmarcAL1 maintains a physiological association with Angptl3 at peroxisomes under normal growth conditions, with the interaction significantly enhanced during slow cell growth.

Biochemical analysis further confirms the shuttling of SmarcAL1 between nucleus and cytoplasm. Under normal growth conditions, most SmarcAL1 molecules were present in nuclei (Fig. 3C, top). During serum starvation, the majority of SmarcAL1 translocated from nuclei to peroxisomes (bottom). Shuttling also occurred in Fc control cells due to endogenous Angptl3, but it was more drastic in Fc-Angptl3 cells. Once again, in Fc- Angptl3 cells with low serum, peroxisomal SmarcAL1 was heavily modified.

Thus, the results from our biochemical and immunofluorescence analyses are highly consistent, indicating that Angptl3 regulates the SmarcAL1 shuttling between nuclei and cytoplasm/peroxisomes in response to cell growth states.

### Angptl3-regulated cellular lipid metabolism is dependent on SmarcAL1 activities

The above results confirmed the cellular lipid phenotypes induced by Angptl3 expression. Previously, we demonstrated that SmarcAL1 regulates the expression of key lipid genes crucial for cellular lipid homeostasis ^21^. The discovery of Angptl3’s role in mediating the cytoplasmic translocation of SmarcAL1 suggests that the lipid phenotypes under Angptl3 expression may be linked to the activity of SmarcAL1 in gene expression. The cell models under the two growth conditions would enable us to test this hypothesis.

We conducted microarray analyses to investigate the gene expression changes caused by Angptl3 overexpression under serum starvation (Fig. 4).

Under normal growth media, Angptl3 expression up-regulated the expression of numerous lipid genes (Fig. 4C). The expression of many genes related to FA metabolism was also up-regulated, driving several cellular lipid catalytic processes. These up- regulated changes may represent cellular adaptation due to stable Angptl3 expression. Interestingly, serum starvation reversed this expression profile by decreasing the expression of genes related to lipid metabolism and FA metabolism, consequently down- regulating various cellular lipid catalytic processes. Conversely, the expression of genes related to insulin resistance increased under serum starvation (Fig. 4D). Common key lipid genes were shared between the two conditions, up-regulated under normal growth and down-regulated under serum starvation (Fig. 4E). It is especially worthwhile to notice that the expression of the genes related to peroxisome activities were up-regulated under normal growth and down-regulated under serum starvation (Fig. 4C and 4D).

We hypothesized that the gene expression changes under serum starvation might result from SmarcAL1’s translocation to the cytoplasm, involving a dose-related allosteric interaction with Angptl3 at peroxisomes. Since we have both ANGPTL3 ^18^ and SMARCAL1 KO ^21^ cells we could test this hypothesis by examining whether the absence of ANGPTL3 leads to increased SMARCAL1 in nucleus (sFig. 6). In our previous report, we showed that, in the SMARCAL1 KO cells, there was minimal SMARCAL1 signal in both the nucleus and cytoplasm ^21^. However, in the ANGPTL3 KO cells, the SMARCAL1 nuclear signal relatively increased compared to that in the nuclei of control cells (sFig. 6). These contrasting results could have different effects on gene expression changes, reflecting SMARCAL1’s transcriptional regulation. To explore this, we compared the RNA-seq data from SMARCAL1 KO and ANGPTL3 KO cells (Fig. 5). Indeed, many genes exhibited increased expression in ANGPTL3 KO cells and decreased expression in SMARCAL1 KO cells, and vice versa. Interestingly, the correlation of the former comparison is much stronger than the latter (r=-0.61 vs r=-0.27, Fig. 5A vs 5B), indicating that those expression changes might be directly related to SMARCAL1 activity. These KO cells did share some common genes with opposing expression patterns, including key genes for cellular lipid metabolism such as PPARG, GPAM, and MPV17L (Fig. 5C).

**Fig. 6.**
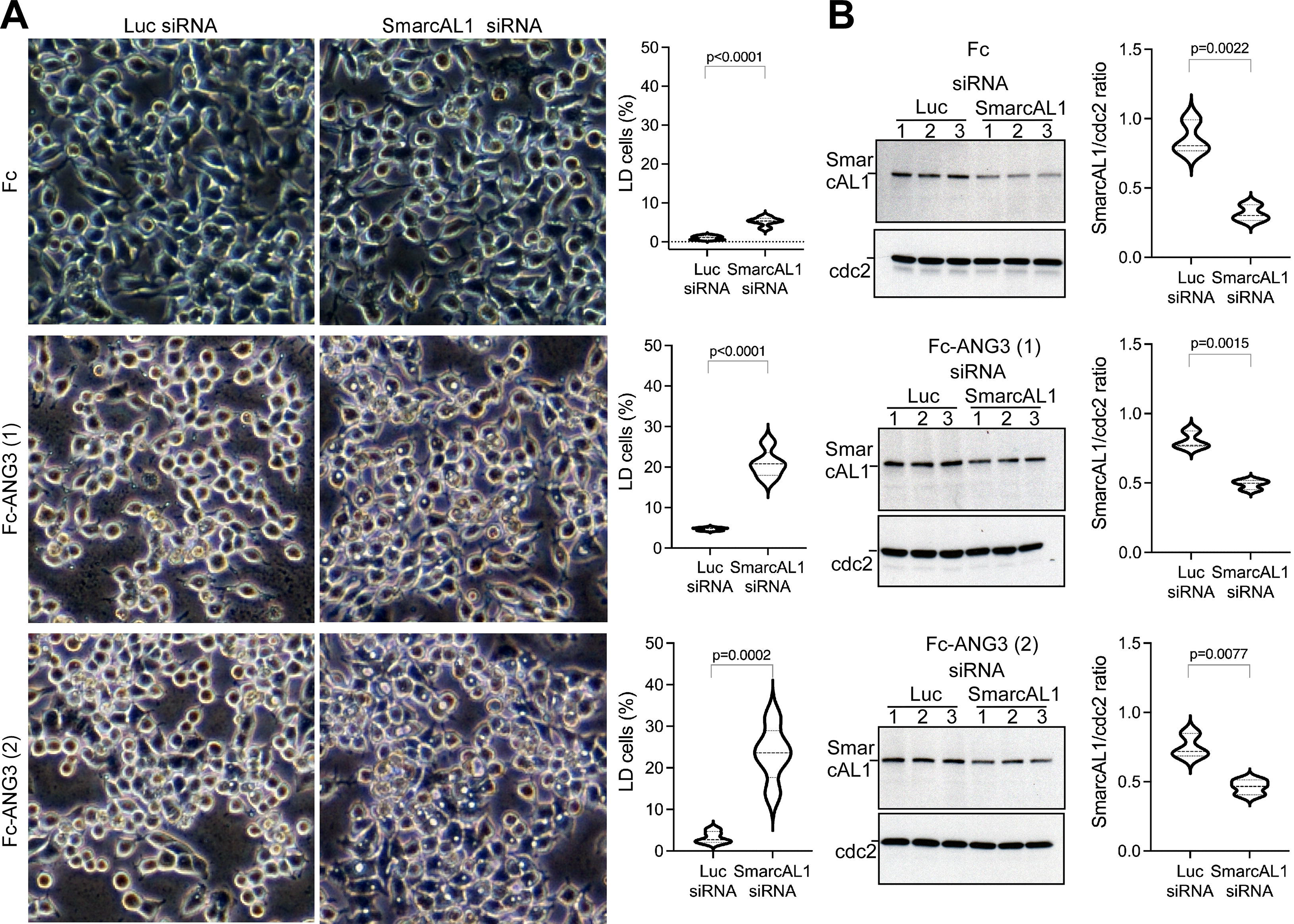
**Silencing SmarcAL1 expression increased LD formation under the normal growth condition. A**. Reduced SmarcAL1 expression increases LD formation at the normal growth condition. Fc and Fc-Angptl3 (with two replicates) cells were transfected with SmarcAL1 siRNA (siRNA1 plus 2) and equivalent luciferase (Luc) siRNAs (three replicates for each). The siRNA-transfected Fc and Fc-Angptl3 cells were grown with DMEM media with 10% FBS at 37 °C. Images were obtained under light microscopy (left panels). Quantification of LD formation (right panels). **B**. Western analysis of SmarcAL1 expression. Extracts from the transfected cells as in A were analyzed with Western blotting and detected with anti-SmarcAL1 and -cdc antibodies (left panels). Quantification of SmarcAL1 expression (right panels). Quantification was based on the ratios of SmarcAL1 expression over cdc expression. The blots were quantified using ImageJ.

To further support the connection, we used siRNAs to downregulate the expression of SmarcAL1 in the stable cell lines utilized in Fig. 1A. In Fc cells, silencing SmarcAL1 led to a slight yet significant increase in the number of LDs. In contrast, the silencing of SmarcAL1 in Fc-Angptl3 cells resulted in a significant enhancement of LD formation (Fig. 6). We reasoned that under increased Angptl3 expression, reduced SmarcAL1 expression would exacerbate LD formation. All these assays were conducted under normal growth conditions. The findings indicate that in the presence of increased Angptl3 expression, decreased SmarcAL1 expression prompts the cells to accumulate fat.

### Angptl3 modulates blood TG levels in a manner associated with its regulation of SmarcAL1 activities

To investigate the in vivo effects of Angptl3 and SmarcAL1 activities on blood TG levels, we conducted functional analyses in mice. Previous studies have examined the effects of modulating Angptl3 expression in mouse models, primarily through virus-mediated gene transfer. These approaches resulted in transient elevations in blood TG levels within 7 days, which gradually declined within 14 days ^1,7^. However, due to the sustained expression of Angptl3 from the virus-mediated approach and a lack of expression data from the studies, it has been challenging to assess the correlation between Angptl3 levels and changes in blood TG. Additionally, several studies utilized purified recombinant Angptl3 proteins injected into mice to evaluate their effects on blood TG levels within 24 hrs ^1,25^. One study demonstrated a transient increase in blood TG levels following protein injection, which returned to baseline levels within 6 hrs.

In this study, we intended to utilize Angptl3 recombinant protein to focus on its longer-term effects on TG levels. We monitored blood TG levels ten days after the initial injection to provide a complementary understanding of Angptl3’s impact on lipid metabolism. The Fc and Fc-Angptl3 proteins, purified from the cells we used in Fig. 1A (sFig. 7), were injected three times (two days in between) into female wild-type (WT) mouse littermates fed a TG-rich high-fat diet (HFD) for 4 weeks. The results demonstrated that Fc-Angptl3, but not Fc, effectively decreased plasma TG levels within ten days post-initial injection (Fig. 7A, left), without apparently influencing TC, HDL-C, and non-HDL-C levels. To validate this observation, we repeated the experiment using male WT mice under identical conditions, and once again, Angptl3 protein was found to reduce blood TG levels in these mice without affecting other lipid levels (Fig. 7A, right).

**Fig. 7.**
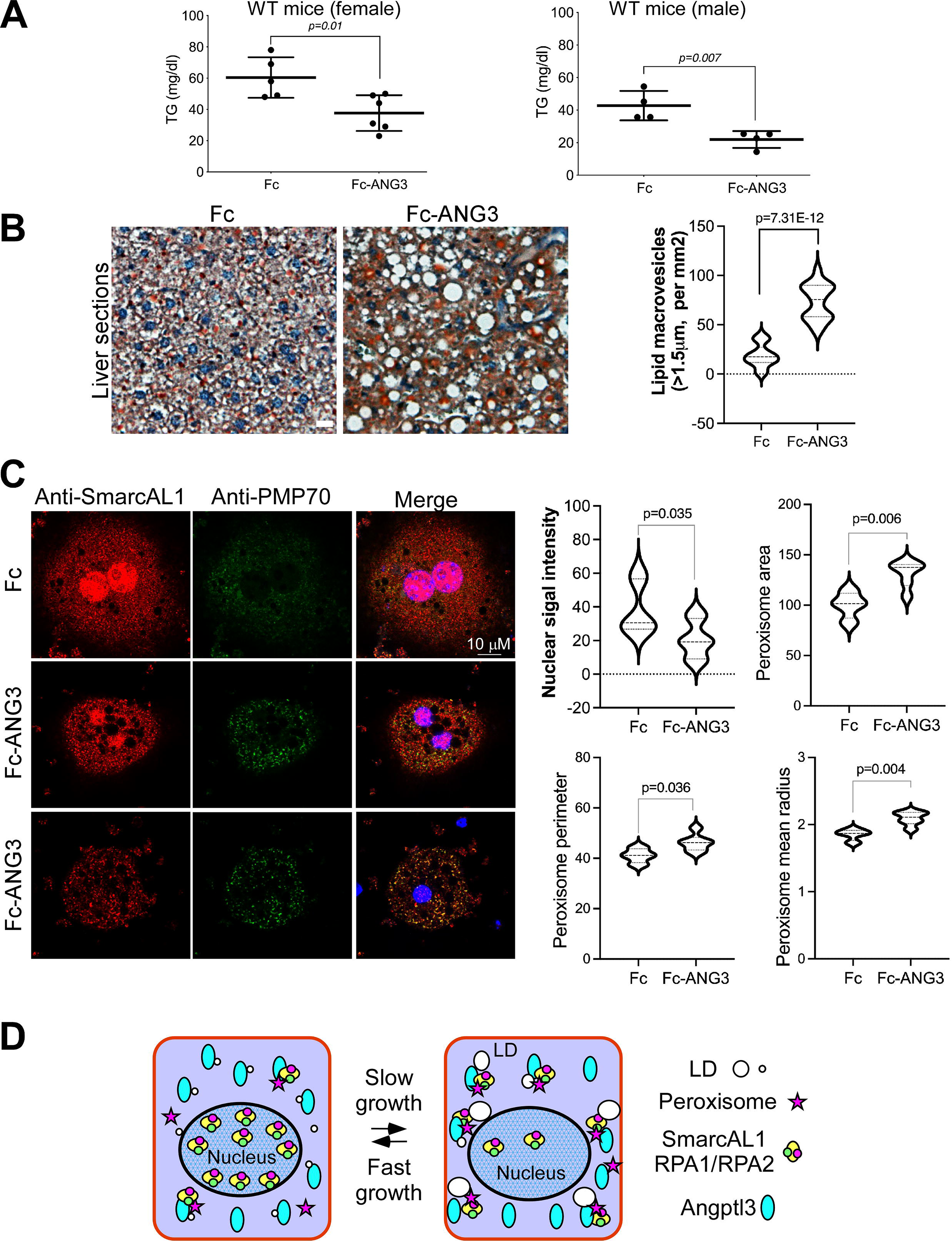
**Angptl3 promotes blood TG clearance that is associated with SmarcAL1 activities**. **A**. Angptl3 protein has a long-term effect on reducing blood TG levels. Female (left) and male (right) WT mice (16 weeks) were fed with HFD for 4 weeks. Purified Fc and Fc-Angptl3 proteins as in sFig. 7 were injected into these mice. Blood samples were collected ten days after the first injection and analyzed for plasma TG levels (see Methods for experimental details). **B**. Angptl3 promotes fat accumulation in the livers. Liver sections from the mice were stained with oil red staining and large lipid macrovesicles were quantified using ImageJ (right). Bar, 25 μM. **C**. Ex vivo analysis of mouse primary hepatocytes. Primary hepatocytes harvested from the livers of the mice as in A (see Methods for details) were fixed and hybridized with anti-PMP70 (green) and -SmarcAL1 (red) plus fluorescence-labeled secondary antibodies. Images were collected from confocal microscopy. Quantification was conducted using CellProfiler. Bar, 10 μM. **D**. Schematic representation of the role of Angptl3-SmarcAL1 interaction in TG storage and trafficking. Under increased Angptl3 expression, slow cell growth induces SmarcAL1 translocation from nucleus to cytoplasmic peroxisomes by interacting with Angptl3 favoring fat storage. Fast cell growth reverses the process, and SmarcAL1 moves back to nucleus for regulating the expression of key lipid genes responsible for lipid metabolism. P values shown in this figure were calculated from two-sided unpaired t-test.

In our cell models, increased Angptl3 expression led to elevated cellular TG accumulation and LD formation. Given that Angptl3 is exclusively expressed in the liver, any effects from the recombinant protein on tissue TG distribution would likely be observed in this organ. Therefore, we harvested the livers from the mice and examined potential changes in TG distribution. We observed that Angptl3 protein injection induced a significant increase in the formation of LDs (∼3.9 times), indicating fat accumulation in the livers (Fig. 7B).

To investigate whether Angptl3-mediated TG clearance is related to SmarcAL1 activities, we conducted ex vivo analyses to assess the regulation (Fig. 7C). Primary hepatocytes were harvested from the livers of mice injected with Fc or Fc-Angptl3 proteins, as described above. Subsequently, hepatocytes were subjected to immunofluorescence analysis similar to that conducted for our cell models as in Fig. 2. In hepatocytes from mice injected with Fc protein, SmarcAL1 was enriched in the nuclei with some background signal in the cytoplasm. Contrastingly, in hepatocytes from mice injected with Fc-Angptl3 protein, the SmarcAL1 signal was weaker in the nuclei and showed enhanced signal from cytoplasmic particles, largely co-localized with the peroxisome marker, PMP70. In some cells, the nuclear SmarcAL1 signal was nearly undetectable. These cells exhibited larger peroxisome particles compared to control cells (Fig. 7C, right). These results suggest that, akin to the cell models, increased Angptl3 induces the translocation of nuclear SmarcAL1 to cytoplasmic peroxisomes, leading to cellular TG accumulation (Fig. 7D). Thus, these results are well consistent with the observation from cell models, suggesting that Angptl3 has an additional role in regulating cellular lipid metabolism in addition to its inhibition of LPL.

## Discussion

In this study, we demonstrated that Angptl3 modulates cellular lipid metabolism. Increased Angptl3 expression induces cellular fat accumulation under conditions of slow cell growth. Our detailed analysis reveals that slow cell growth enhances the interaction of Angptl3 with SmarcAL1, specifically at the cytoplasmic peroxisomes. This interaction induces expression changes in numerous lipid genes, facilitated by SmarcAL1 translocation, promoting the TG accumulation. Consequently, our study underscores that the chromatin regulator SmarcAL1 serves as the cellular target of Angptl3, and the interactions between Angptl3 and SmarcAL1 may present an additional mechanism that enables cells to modulate TG storage and trafficking in response to the cell growth states. The findings presented in this study offer additional insights into the role of Angptl3 in regulating lipid metabolism. Intravascular lipase inhibitions have long been thought the major mechanism of ANGPTL3, explaining most lipid phenotypes observed in individuals carrying ANGPTL3 mutations ^10,11,26^. While this mechanism aligns well with low blood TG levels in individuals with ANGPTL3 loss of function (LoF) mutations due to LPL inhibition ^3^, it does not fully account for all observed lipid phenotypes. For instance, individuals with ANGPTL3 LoF mutations also exhibit low blood FA levels ^27^. Consistently, Angptl3 KO mice also exhibit low blood FA levels ^28^. Notably, injecting Angptl3 proteins into mice concurrently raised blood TGs and FAs ^1^ (discussed below). Although there are alternative mechanisms to explain these discrepancies ^12,13^, the changes in FA levels do not directly correlate with low or high LPL activities, with or without inhibition by ANGPTL3. These disparities suggest additional insights into Angptl3’s role in lipid metabolism.

Our study sheds light on the role of Angptl3 in regulating cellular lipid metabolism, a notion that has been suggested in earlier research ^12,13^ and is increasingly supported by recent investigations. We previously demonstrated that siRNA-mediated silencing of ANGPTL3 induced the combined hypolipidemia in mice, with lower LDL-C levels likely resulting from reduced nascent ApoB-100 (or VLDL) secretion, while ANGPTL3 gene deletion caused TG accumulation in human cells ^18^. These findings are in line with recent independent studies. For instance, ANGPTL3 deficiency in HepG2 cells led to ∼50% reduction in nascent ApoB-100 secretion due to early lipoprotein assembly defects ^19^. siRNA-mediated ANGPTL3 silencing increased cellular fat accumulation in human cells, likely due to decreased energy consumption ^20^. Notably, evidence from clinical trials targeting ANGPTL3 further underscores the importance of proper Angptl3 expression in cellular lipid metabolic processes. ASO-based silencing of ANGPTL3 in statin-treated patients with elevated cholesterol resulted in dose-dependent hepatic fat accumulation ^17^, whereas another clinical trial utilizing monoclonal antibodies targeting ANGPTL3 did not report similar side effects ^29^. These studies provide direct evidence of the critical role of Angptl3 expression in regulating cellular lipid metabolism.

Our proteomics and functional analysis revealed a direct interaction between Angptl3 and SmarcAL1, demonstrating that SmarcAL1 is the primary cellular target of Angptl3. Angptl3 was proposed as a hormone that regulates the energy trafficking, storage, and expenditure ^30^. This interaction raises a possibility that SmarcAL1 could serve as the intracellular receptor of Angptl3, analogous to the interaction between angiopoietins and their corresponding receptors, given the fact that angiopoietin-like proteins share high homology and nearly indistinguishable structure with angiopoietins ^5,31^. The specific interaction with SmarcAL1 suggests that Angptl3 may plays unique roles in cellular lipid metabolism compared to other angiopoietin-like members linked to lipid metabolism, including Angptl4 and Angptl8 as demonstrated in our previous study ^21^.

Our studies reveal that Angptl3 regulates the activities of SmarcAL1 associated with its cytoplasmic translocation, likely underlying the lipid phenotypes related to Angptl3 expression. We previously showed that SmarcAL1 is crucial for cellular lipid homeostasis, largely related to its activities in regulating gene expression ^21^. In this study, we provided two aspects of evidence supporting Angptl3’s regulation of SmarcAL1 activities based on rat and human cell models. First, under increased Angptl3 expression, serum starvation induces the cytoplasmic translocation of SmarcAL1 from nuclei, resulting in decreased expression of many lipid genes and massive LD formation. Second, inactivation of the ANGPTL3 gene in human cells increases the presence of SMARCAL1 in nuclei, resulting in increased expression of many lipid genes. The expression of these genes is down- regulated in SMARCAL1 KO cells, indicating that the changes are likely due to SMARCAL1 activities. Additionally, we confirm the presence of SmarcAL1 at cytoplasmic peroxisomes, and our study demonstrates the dynamic interaction of Angptl3-SmarcAL1 at peroxisomes in response to changes in cell growth state. Given the central role of peroxisomes in FA catalysis ^32^, this finding adds another layer to their regulation of cellular lipid metabolism. Further studies should shed more insights for this regulation.

It is intriguing that the interaction between Angptl3 and SmarcAL1 occurs at cytoplasmic peroxisomes, especially considering that neither protein contains signals that typically target them to peroxisomes. We hypothesize that the localization of SmarcAL1 to peroxisomes may depend on Angptl3. Angptl3 possesses a unique N-terminal coiled-coil (or leucine zipper) domain known to mediate protein-protein interactions ^33^. This domain is notably rich in amino acids such as leucine, lysine, and arginine, which promote the formation of alpha-helix structures ^5,8,34^. Consequently, the abundance of these basic amino acids renders this domain amphipathic, possessing both hydrophilic and lipophilic properties conducive to membrane penetration ^35^. Indeed, coiled-coil proteins have been demonstrated to play critical roles in facilitating various membrane-penetrating events ^33^, suggesting that this property may enable the Angptl3-SmarcAL1 complex to localize to the membrane of peroxisomes. However, further research is necessary to substantiate this hypothesis.

It is likely that TG is the major energy substrate regulated by Angptl3-SmarcAL1 interaction. The findings that Angptl3-regulated peroxisomal translocation of SmarcAL1 and resulting gene expression changes may have broad significance in systematic TG storage and trafficking. Our data indicate that this translocation is related to cell growth states. Under conditions of slow cell growth, the majority of nuclear SmarcAL1 translocates to peroxisomes, where it interacts with Angptl3. The dosage change of SmarcAL1 in nuclei affects the expression of many genes involved in cellular lipid metabolism, resulting in fat accumulation inside cells. This mechanism may provide a regulatory control of systematic energy storage and trafficking by enabling periphery tissues, such as adipose tissues, to respond to feeding states (Fig. 7D). More in vivo analyses are needed to prove this hypothesis.

It is plausible that Angptl3 might exert dual-phase effects on blood TG levels, reflecting its intravascular and cellular functionalities. Previous investigations leveraging Angptl3 recombinant proteins for functional analyses have revealed a transient increase in blood TG levels in mouse models ^1,7^. For instance, one study showed a rapid elevation within 2 hrs post-Angptl3 protein injection, followed by a swift decline within 6 hrs, primarily attributed to LPL inhibition, as proposed in the study. To explore the prolonged effects of Angptl3, we evaluated TG levels ten days post-protein injection. Intriguingly, we observed a decrease in TG levels concurrent with increased fat accumulation in the livers. The decrease may imply a potential long-term effect on blood TG independent of LPL inhibition. Indeed, ex vivo analysis of primary hepatocytes from these mice confirmed that Angptl3 protein induced the translocation of SmarcAL1 from nuclei to cytoplasmic peroxisomes. These observations correlate well with in vitro data presented in our study, demonstrating that increased Angptl3 expression induces cellular TG accumulation. This suggests that Angptl3, through its interaction with SmarcAL1, plays an additional role in regulating lipid metabolism.

While our study has contributed valuable evidence supporting the additional role of ANGPTL3 in cellular lipid metabolism regulation, it is important to acknowledge certain limitations when interpreting our findings. We conducted detailed analyses to investigate how changes in Angptl3 expression induce cellular lipid phenotypes and regulate the activities of SmarcAL1 in cellular metabolism. However, it is notable that the majority of our results are derived from cell models, and the in vivo evidence is limited. Future research endeavors should aim to encompass broader in vivo analyses utilizing diverse mouse models. Additionally, tissue-specific knockout studies targeting SmarcAL1 in mice would offer deeper insights into the Angptl3-SmarcAL1 interaction and its role in regulating energy storage and trafficking.

In our previous study, we showed that the genetic variations at SMARCAL1 locus are associated with body mass index and other metabolic phenotypes ^21^. To better understand the clinical relevance of our findings, it is important to investigate whether and how these phenotypes are related to the interaction of SMARCAL1 with ANGPTL3. Such studies could involve analyzing patient data, or functional analyses of ANGPTL3-SMARCAL1 interaction using patient-derived cell models. Moreover, further efforts should be also directed towards unraveling the mechanistic insight about ANGPTL3-SMARCAL1 interaction on peroxisome and how this interaction is related to the peroxisome function in regulating the FA catalytic process. These future studies would enhance our understanding of the physiological relevance of the Angptl3-SmarcAL1 axis in human lipid metabolism regulation.

## Materials and Methods

### Antibodies and biological reagents

All antibodies and reagents utilized in this study are listed in Supplemental Table 3.

### Cell culture, plasmid constructs, SmarcAL1 siRNAs, and transfection

McA, HeLa, Huh7, and HepG2 cells, as well as the cell lines generated in this study, were cultured in DMEM medium (Invitrogen) supplemented with 10% FBS (Sigma) or OPTI-MEM media with 2% low-IgG FBS, as specified in individual figures. For purification purposes, low-IgG FBS (Fisher Scientific) was utilized for cell culture. The SMARCAL1 KO and ANGPTL3 KO and respective control Huh7 cells were generated as previously described ^18,21^.

The Fc and Fc-Angptl3 expression constructs and cell lines were generated as previously described ^21^. The GFP-Angptl3 construct was generated by cloning the cDNA of human ANGPTL3 into the pEGFP-C2 vector (Clontech) at EcoRI and ApaI sites. Fc- or GFP-tagged N-terminal Angptl3 constructs were created by cloning the first half (690 bp) of human ANGPTL3 cDNA into the respective plasmids mentioned above. Inducible Fc and Fc-Angptl3 expression constructs were built using the Tet-on expression system (Clontech). The cDNA of Fc was cloned into the pTRE-HA vector at BamHI-HindIII sites, followed by insertion of human ANGPTL3 cDNA into the Fc-pTRE-HA vector at HindIII-NotI sites. In both constructs, the HA tag was replaced with the Fc tag. All constructs underwent thorough validation through DNA sequencing to ensure accuracy and fidelity.

Transfection was carried out using Lipofectamine 2000 (Invitrogen) in accordance with the manufacturer’s instructions. Stable Fc and Fc-Angptl3 McA cell lines were as previously described ^21^. HepG2 tet-on cells were produced by transfecting pTet-on plasmids, followed by selection with G418. Inducible Fc and Fc-Angptl3 cells were obtained by transfecting the inducible constructs into HepG2 tet-on cells, followed by hygromycin selection. For stable and inducible Fc and Fc-Angptl3 cells, single clones were isolated by seeding single cells in 96-well plates and subsequently screened using Western blotting analyses. GFP, GFP-N-Angptl3, and GFP-Angptl3 expression constructs were utilized for transient transfection.

Two siRNAs, siRNA1 (5’-GCUGGUGGUUGUGCCUUCC-3’) and siRNA2 (5’-ACAUACGGUGGCAGUGUUG-3’), targeting rat SmarcAL1 mRNA, along with a control luciferase siRNA (5’-GCCAUUCUAUCCUCUAGAGGAUG-3’), were procured from Qiagen. Transfection of the siRNAs into cells (in triplicate) was conducted using RNAiMAX (Invitrogen) following the manufacturer’s instructions. The transfected cells were then utilized for phase imaging and Western blotting analyses, as described in Fig. 6. Cell duplicates were treated under identical conditions.

### Cell extracts, peroxisome-enriched fraction, and Western blotting analysis

Whole cell extracts were prepared by lysing cells in a buffer containing 150 mM NaCl, 50 mM Tris (pH 7.5), 1% IGPAL-CA-630 (Sigma # I8896), and a protease inhibitor cocktail (Roche), followed by rotation at 4°C for 15 mins and centrifugation at 13,000 rpm for 10 mins. For the preparation of the peroxisome-enriched fraction, cells were washed twice with ice-cold PBS and then twice with ice-cold HES buffer composed of 5 mM HEPES (pH 7.4), 0.5 mM EDTA, 250 mM sucrose, and a complete protease inhibitor cocktail. Subsequently, cells were homogenized by passing through a 25-gauge needle ten times, followed by centrifugation at 1,000 g for 10 mins at 4°C to pellet nuclei. The peroxisome-enriched fraction was obtained by centrifuging the post-nuclear supernatant at 16,000g for 20 mins at 4°C ^36^. Western blotting analyses were conducted using precast Mini-PROTEAN TGX SDS-PAGE gels (4-20%) (Bio-Rad). Blots were probed with primary antibodies as indicated in the figures and appropriate secondary antibodies.

### Affinity pull-down (PD) assay, and recombinant protein production and purification

Fc and Fc-Angptl3 affinity PD assays were conducted following established protocols ^37,38^. Briefly, protein-A beads were washed with lysis buffer, followed by incubation with cell extracts, rotating at 4°C for ∼12 hrs. The mixtures were then centrifuged at 2,000 rpm for 5 mins, and the beads were washed three times with lysis buffer. Subsequently, the precipitates were loaded onto precast Mini-PROTEAN TGX SDS-PAGE gels (4-20%) for Western blot analysis.

Recombinant Fc and Fc-Angptl3 proteins were purified from the OPTI-MEM culture media with 2% low-IgG FBS obtained from Fc and Fc-Angptl3 McA cells. The media was dialyzed against 1XTBS, followed by centrifugation at 8,000 rpm for 30 mins at 4°C and filtration. Next, the media was incubated with protein-A beads, rotating for ∼12 hrs at 4°C, and then centrifuged at 2,000 rpm for 20 mins at 4°C. The precipitates were washed with 1XTBS and eluted with 0.3 M glycine (pH 3). Following neutralization, eluates were dialyzed against PBS. After filtration, protein concentration was measured before use.

### RNA extraction, microarray and RNA-seq transcriptome-wide analyses

Total RNA was isolated using the RNeasy Mini Kit (Qiagen) following the manufacturer’s instructions. Purified RNA samples underwent Bioanalyzer trace analysis (Agilent) to assess RNA quality. All RNA samples (each with three replicates) utilized for transcriptome analyses met optimal specifications for total RNA, including a RIN score of > 7.0, and RNA purity ratios of 260/280 = 1.8-2.0 and 260/230 = 1.8-2.0, which are key markers of RNA integrity. Microarray and RNA-seq analyses were conducted at the Mass General Brigham Biobank Genomics Core. Microarray assays were performed using Affymetrix GeneChip Rat Gene ST arrays (for McA cells) or GeneChip Human Gene 2.0 ST arrays (for Huh7 cells) (Applied Biosystem/Thermo Fisher). GeneChips were scanned using the Affymetrix GeneChip Scanner 3000 7G running Affymetrix Gene Command Console ver 3.2. Microarray data were analyzed using Affymetrix Expression Console (ver 1.3.0.187) using the analysis algorithm RMA. For RNA-seq analyses, libraries were generated using TruSeq® Stranded Total RNA Library Prep Human/Mouse/Rat (20020597, Illumina). Sequencing reads with 75 bp paired ends were generated on an Illumina HiSeq 2500 platform following the manufacturer’s protocol. Raw reads were processed, and trimmed reads were aligned to the human genome (hg19) or rat genome (Rattus_norvegicus.Rnor_6.0.85.chr.gtf) using the STAR aligner (2.4.2). Subsequent differential expression analysis was performed using cuffdiff in cufflinks (2.2.1). Differentially expressed genes with p and q values less than 0.05 were subjected to gene ontology (GO) enrichment analysis using DAVID (the database for annotation, visualization, and integrated discovery) (https://david.ncifcrf.gov) ^39^. Pathways and biological processes with Bonferroni, Benjamini, and false discovery rate (FDR) less than 0.05 were considered statistically significant.

### Immunofluorescence and imaging analyses

All phase images were captured using a standard light microscope. Immunofluorescence analysis was conducted essentially as previously described ^40^. Briefly, cells were fixed with a PBS buffer containing 3% paraformaldehyde (Sigma) and 2% sucrose for 10 mins, followed by permeabilization with Triton X-100 buffer (0.5% Triton X-100, 20 mM HEPES-KOH, pH 7.9, 50 mM NaCl, 3 mM MgCl2, 300 mM sucrose).

Subsequently, the cells were blocked in PBS buffer containing 0.5% bovine serum albumin and 0.2% gelatin (Sigma) before incubating with primary antibodies and appropriate secondary antibodies. Coverslips were mounted in mounting media (Vector Laboratories). All fluorescence images were acquired by scanning the slides using a Leica SP5 AOBS Scanning Laser Confocal Microscope. Image analysis was performed using CellProfiler or as otherwise described in individual figures. Consistent thresholds were applied to all images in the relevant series of experiments, and particle analysis was utilized to collect cell particle number, size and intensity data.

### Mice, mouse treatments and handling

Wild-type mice (C57BL/6J) were purchased from Jackson Laboratory. All mice were maintained on a standard chow diet and housed under a 12-hour light/12-hour dark cycle. All animal procedures were conducted in accordance with the guidelines approved by the Institutional Animal Care and Use Committee (IACUC) and complied with relevant local, state, and federal regulations.

For the protein injection experiments, both female and male WT mouse littermates were randomly divided into two groups: one receiving Fc protein injection and the other receiving Fc-Angptl3 protein injection. Prior to protein injections, the mice were fed a TG-rich high-fat diet (Envigo) for 4 weeks. Purified recombinant proteins were administered via retro-orbital sinus injection post-anesthesia three times (∼25 μg each injection), with 2 days in-between injections.

To minimize variations in blood TG levels, mice were subjected to a fasting and refeeding regimen as previously described ^41^. Briefly, mice underwent fasting for 12 hours followed by refeeding for 12 hours for three consecutive days prior to blood collections. At the conclusion of the fasting period (ten days after the first injection), blood samples were collected from the retro-orbital sinus following anesthesia procedures and immediately centrifuged to prepare plasma. After blood collections, mice underwent primary hepatocyte isolation treatment as described below.

Measurement of mouse plasma TG levels was conducted using a triglyceride colorimetric assay kit (Cayman) according to the manufacturer’s protocol. Additionally, total cholesterol (TC), and high-density lipoprotein cholesterol (HDL-C) levels were enzymatically measured in individual mouse plasma samples using an ACE Axcel chemistry analyzer (Alfa Wassermann). Non-HDL-C levels were estimated indirectly by subtracting the HDL-C from the TC.

### Primary mouse hepatocyte solation and culture

Primary mouse hepatocytes were isolated and cultured following established protocols^42^. Briefly, mice were anesthetized, and abdominal surgery was performed to expose the visceral vena cava without causing damage to blood vessels. A catheter was inserted into the vena cava via the right atrium for perfusion. Perfusion was initiated with Solution 1 containing Hank’s Balanced Salt Solution (HBSS, Gibco) supplemented with 0.5 mM EDTA (pH 8) at a flow rate of 5 mL/min for 5-7 mins. This was followed by perfusion with Solution 2 consisting of DMEM supplemented with collagenase type I (0.8 mg/mL) (VWR International) for 7–8 mins at the same flow rate. Following perfusion, the livers were excised and placed in tubes containing Solution 2. Subsequently, the livers were sliced to release the primary hepatocytes. The isolated hepatocytes were then centrifuged and washed twice with DMEM media. Cell counting and viability assessments were performed, and the hepatocytes were cultured in DMEM supplemented with 2% BSA for subsequent analyses.

### Total cell lipid and polar lipid extract preparation and metabolite mass spectrometry analysis

The metabolomics analysis was conducted at the Metabolomics Program, Broad Institute of MIT and Harvard. Preparation of total cell lipid and polar lipid extracts was performed according to established protocols ^43^. Briefly, for lipid extraction, cells were rinsed with cold PBS (no Mg^2+^/no Ca^2+^), and then 800µL of ice-cold isopropanol (HPLC grade) was added immediately. The cells were scraped and transferred to a 1.5 mL tube, then kept at 4°C for 1 hr. For polar extraction, after the wash step, 800µL of -80 °C 80% methanol (LC/MS grade) was added to the cells, followed by transfer to -80°C for 15 mins. Subsequently, the cells were scraped and collected into a 1.5 mL tube. For both extracts, the cell suspension was vortexed, and cell debris was removed by centrifugation at 9,000 x g, 4°C, for 10 mins. The resulting supernatant was utilized for metabolite mass spectrometry analysis following previously described procedures ^43^. For both assays, three replicates were included for each cell line for testing and cell counting.

### Statistical analysis

Statistical comparisons of groups, including the student’s t-test, were conducted using Prism 10 (GraphPad). Two-way analysis of variance (ANOVA) was employed to compare the matched means of three replicates from experimental groups with those from the control group, as indicated in the figures. Pearson correlation analysis was performed using R by measuring a linear dependence between two variables from every pair of the datasets used, and p-values were calculated with two tails. Detailed group comparisons were described in individual figure legends. In general, p-values of ≤ 0.05 were considered statistically significant (*, p < 0.05; **, p < 0.01; ***, p < 0.001; and ****, p < 0.0001).

## Data deposit

The microarray and RNA-seq data presented in this paper were deposited in Gene Expression Omnibus (GEO). The accession number for the microarray data is GSE221049 (GSE221047 and GSE221048 for the McA cells under normal growth and low serum growth, respectively). The accession numbers of the RNA-seq data from the Huh7 cells with ANGPTL3 KO and SMARCAL1 KO are GSE221371 and GSE221372, respectively.

## Acknowledgements

The authors sincerely thank Dr. Sekar Kathiresan for support to this project. This work was supported by NIH grant, R33HL120781 (S.K.). Y.-X.X. was recipient of Scientist Development Grant (SDG) (11SDG7670007) from the American Heart Association. We thank Richard Lehner (Univ. of Alberta) for McA cell growth and transfection. We thank Kiran Musunuru, Nicolas Kuperwasser, and Chi Gao (MGH) for help at the early stage of this project. We thank Partners Healthcare Personalized Medicine for microarray and RNA-seq analysis, in particular Sami Samir Amr and Mark J. Bowser for their support in the data analysis and submission. We thank Dan Rader and Debra Cromley (Upenn) for mouse blood lipid analysis. We thank Suzanne L. White for mouse tissue processing and oil-red staining and Lay-Hong Ann for the mouse tissue immunohistochemistry and image analysis at Beth Israel Deaconess Medical Center.

## Declaration of interests

The authors declare no competing interests.

## Supplemental figure legends

**sFig. 1. Transient Angptl3 expression increases cellular LD formation. A**. Transient expression of Fc or Fc-Angptl3 in McA cells. **B**. Transient expression of GFP, GFP-N- Angptl3, or GFP-Angptl3 in McA cells. Western analysis of GFP-N-Angptl3 (lane 1) and GFP-Angptl3 (lane 2) expression with anti-GFP antibodies (bottom panel). **C**. Transient expression of Fc or Fc-N-terminal Angptl3 (Fc-N-Angptl3) in HeLa cells. In each panel, the transfected cells were labeled with BODIPY 558/568 C12. The cells were either stained with FITC-labeled anti-Fc antibodies (A and C) or directly visualized for GFP expression (B). Images were obtained from confocal microscopy.

**sFig. 2. Comparative analysis of cellular lipid metabolite changes induced by slow cell growth state**. Total lipid and polar lipid extracts from the cells in media with 10% and 2% FBS were measured with mass spectrometry for metabolite changes. The ratios were calculated by comparing the metabolites from Fc-Angptl3 cells with those from Fc control cells (see sTable 1 for metabolite details). P values for each metabolite group were calculated with two-sided unpaired t-test. P values from two-way ANOVA are indicated.

**sFig. 3. Cellular lipid metabolite profile changes induced by inducible expression of Fc-Angptl3**. **A**. Western analysis of Fc and Fc-Angptl3 expression in HepG2 tet-on cells (with three replicates) treated with or without doxycycline (Dox) as indicated. Western blots were probed with HRP-conjugated antibodies against Fc and β-actin. **B to E**. Total lipid and polar lipid extracts from Fc-Angptl3 HepG2 cells (with three replicates) as in A were measured with mass spectrometry for metabolite profile changes. The ratios were calculated by comparing the metabolites with the averages from those from the control cells. P values from two-way ANOVA are indicated. **F**. Confirmation of Angptl3 interaction with SmarcAL1 in HepG2 tet-on cells. Extracts from Fc and Fc-Angptl3 HepG2 tet-on cells treated with or without Dox were used for expression analysis (top) and for affinity PD assays with protein-A beads. The blots were detected with HRP-conjugated anti-Fc (top) or anti-SmarcAL1 and -Rpa1 (bottom) antibodies.

**sFig. 4. A time course experiment for monitoring real-time SmarcAL1 translocation from nuclei to cytoplasm**. **A**. and **B**. Fc (A) and Fc-Angptl3 (B) McA cells were grown with normal media with 10% FBS (top), and then switched to lower serum media (2% FBS) media for a time course assay. At the indicated time points, cells were fixed and hybridized with anti-PMP70 and -SmarcAL1 plus fluorescence-labeled secondary antibodies. Images were obtained with confocal microscopy. **B**. Quantification of cytoplasmic SmarcAL1 localization. Quantification was carried out from three independent experiments using CellProfiler. P values were calculated with two-sided unpaired t-test or two-way ANOVA as indicated.

**sFig. 5. Low-temperature cell growth under Angptl3 expression accelerates SmarcAL1 cytoplasmic translocation.** Fc (A) and Fc-Angptl3 (B) McA cells were grown in normal media with 10% FBS (top) or in low serum media with 2% FBS (bottom) at 30 °C for three days. The cells were then fixed and hybridized with anti-PMP70, -SmarcAL1 plus fluorescence-labeled secondary antibodies. Images were obtained with confocal microscopy. **C**. Quantification of cytoplasmic PMP70 and SmarcAL1 localization using CellProfiler. P values were calculated with two-sided unpaired t-test as indicated.

**sFig. 6. Comparison of nuclear SMARCAL1 signal between *ANGPTL3* KO and WT Huh7 cells. A**. *ANGPTL3* KO and WT control Huh7 cells under normal growth with 10% FBS media were fixed and hybridized with anti-PMP70, -SmarcAL1 and fluorescence-labeled secondary antibodies. Images were obtained with confocal microscopy. **B**. Quantification of the nuclear signal of SMARCAL1 in *ANGPTL3* KO and WT control Huh7 cells. Quantification was conducted based on three independent experiments using CellProfiler. P values were calculated with two-sided unpaired t-test.

**sFig. 7. Profile of the purified recombinant Fc and Fc-Angptl3 proteins.** The recombinant Fc and Fc-Angptl3 proteins were purified from media of the cells as Fig. 1A and analyzed with Coomassie staining (A) and Western blotting with HRP-conjugated anti-Fc antibody (B).

## Notes

### Competing Interest Statement

The authors have declared no competing interest.

