## Supplementary figures and images for "ANGPTL3 regulates the peroxisomal translocation of SmarcAL1 in response to cell growth states"

### Supplemental Fig. 1

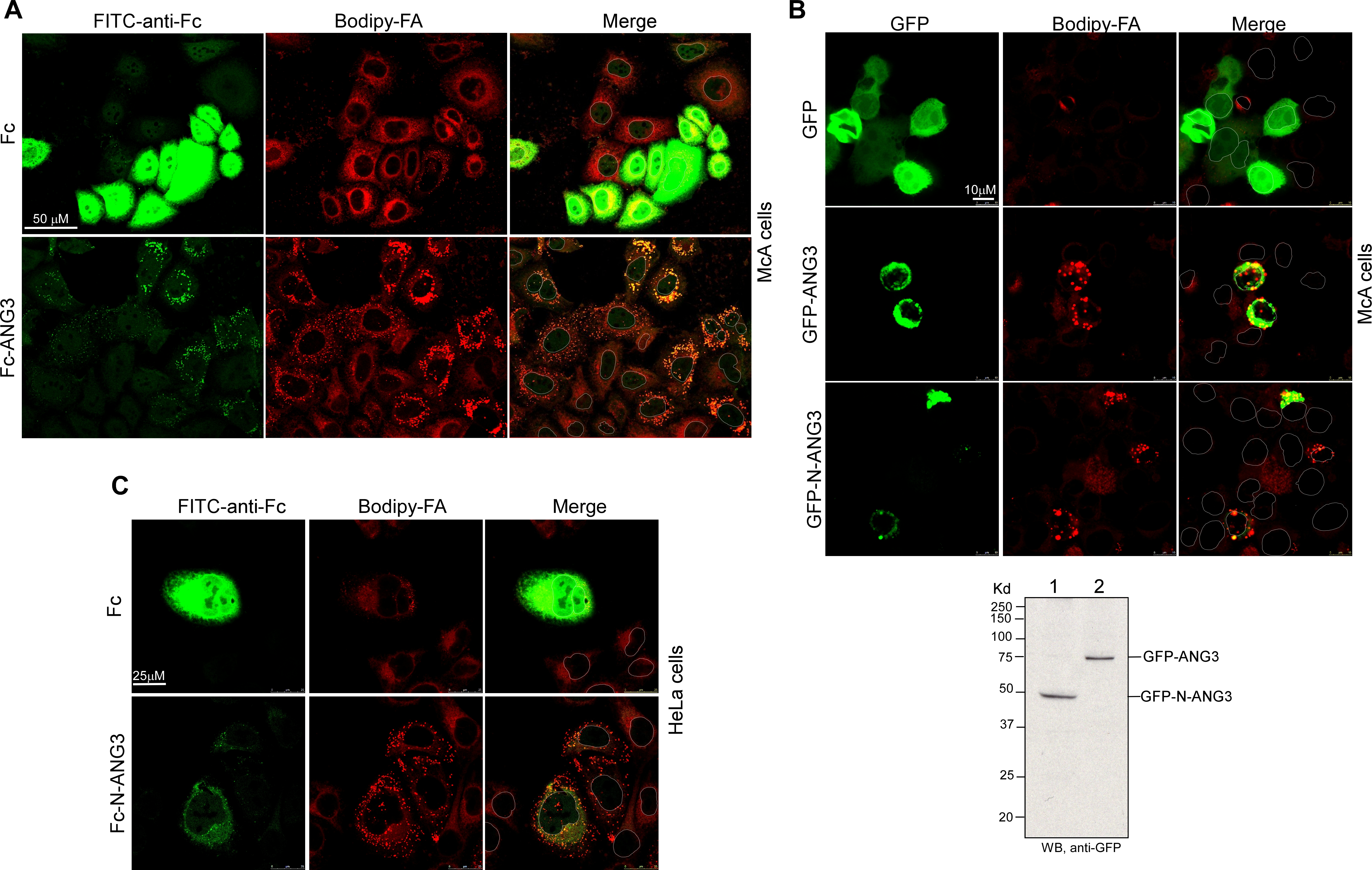

### Supplemental Fig. 2

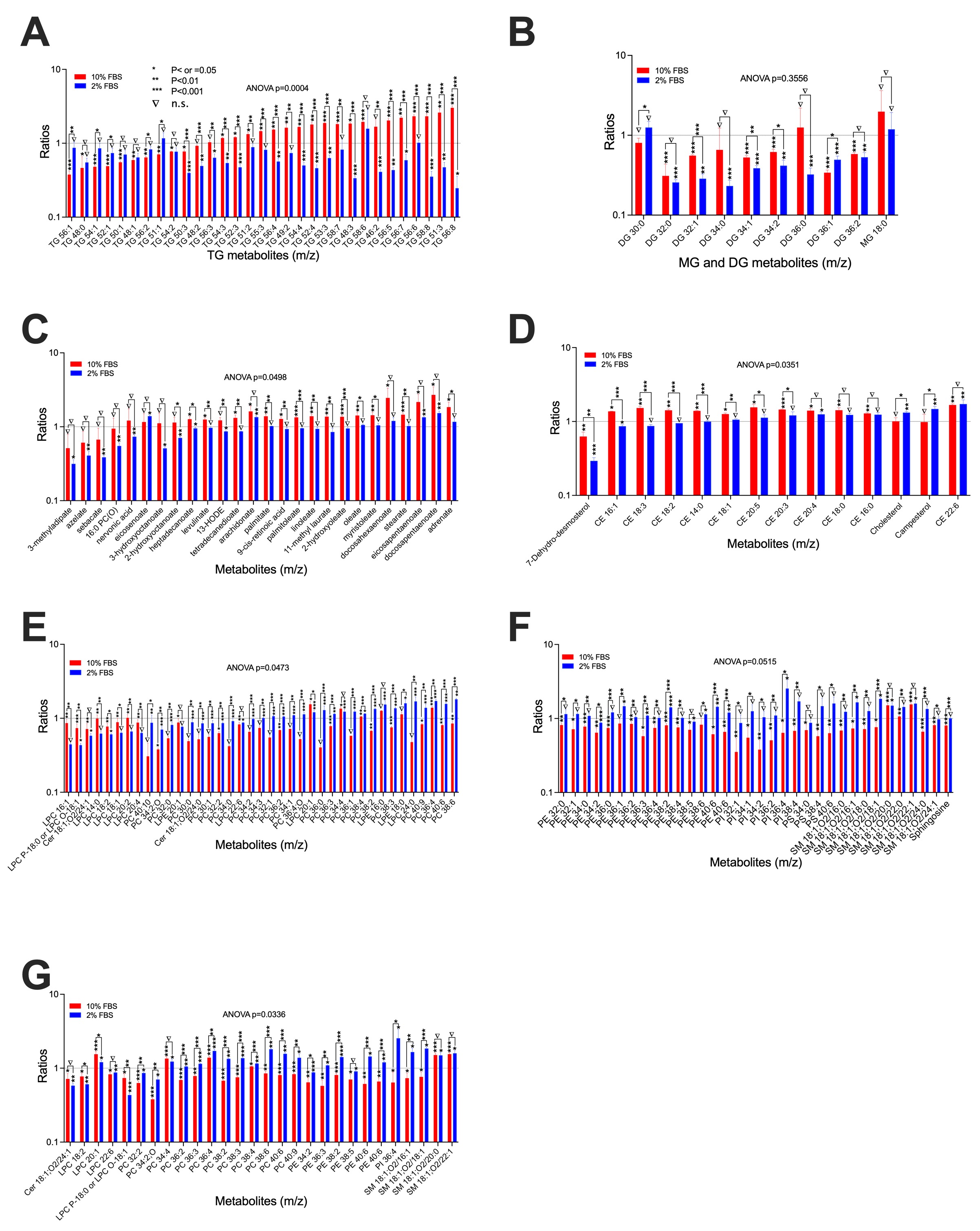

### Supplemental Fig. 3

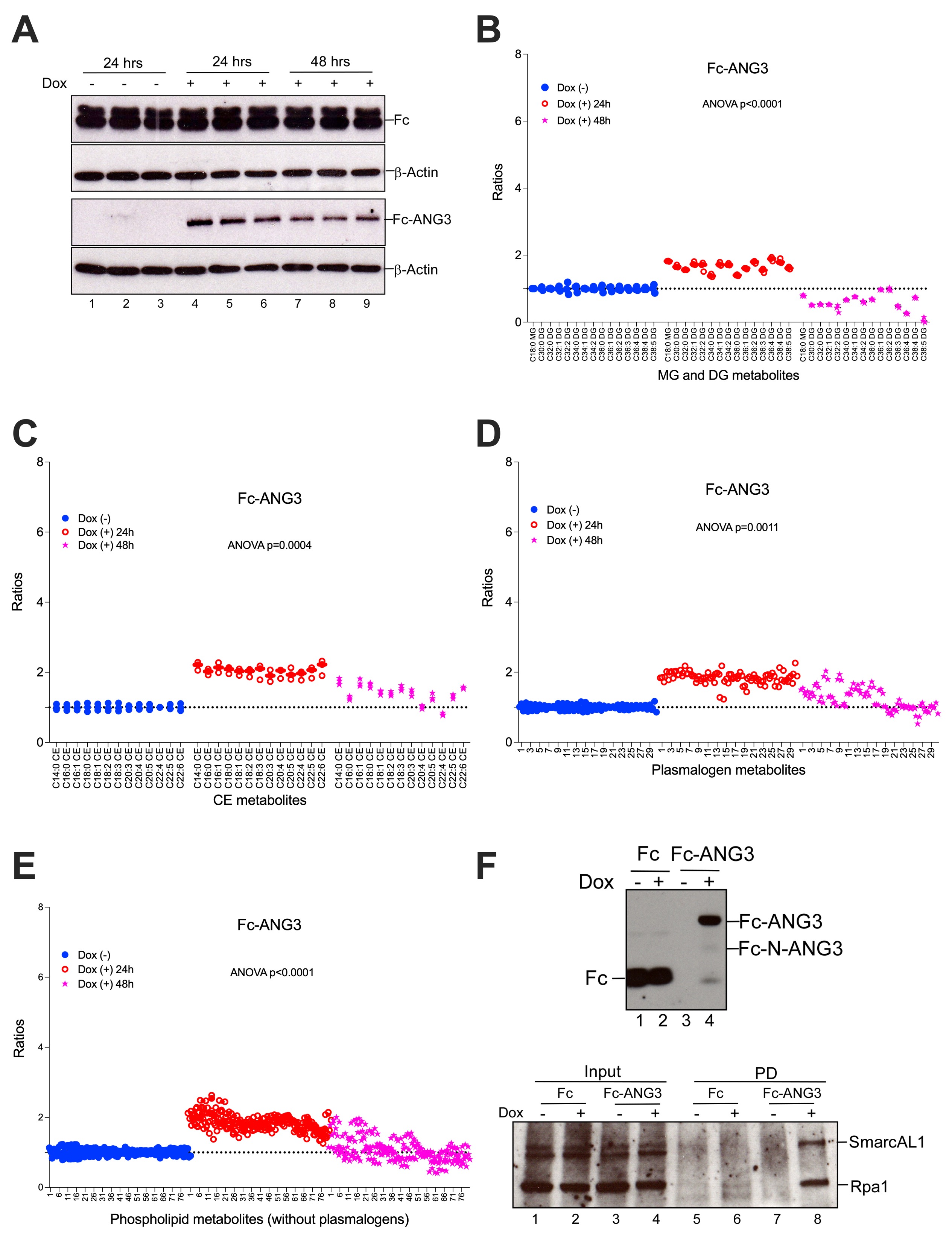

### Supplemental Fig. 4

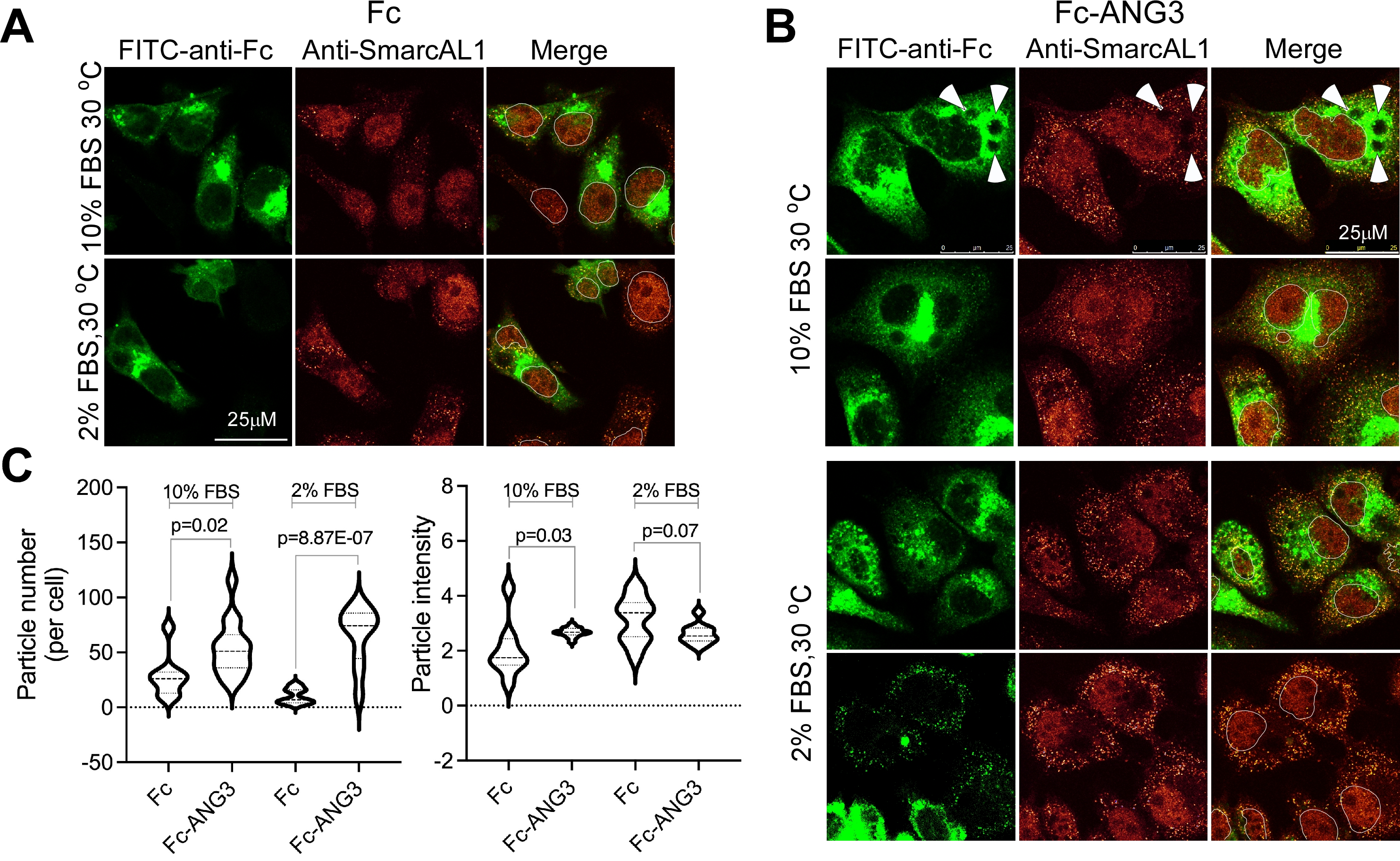

### Supplemental Fig. 5

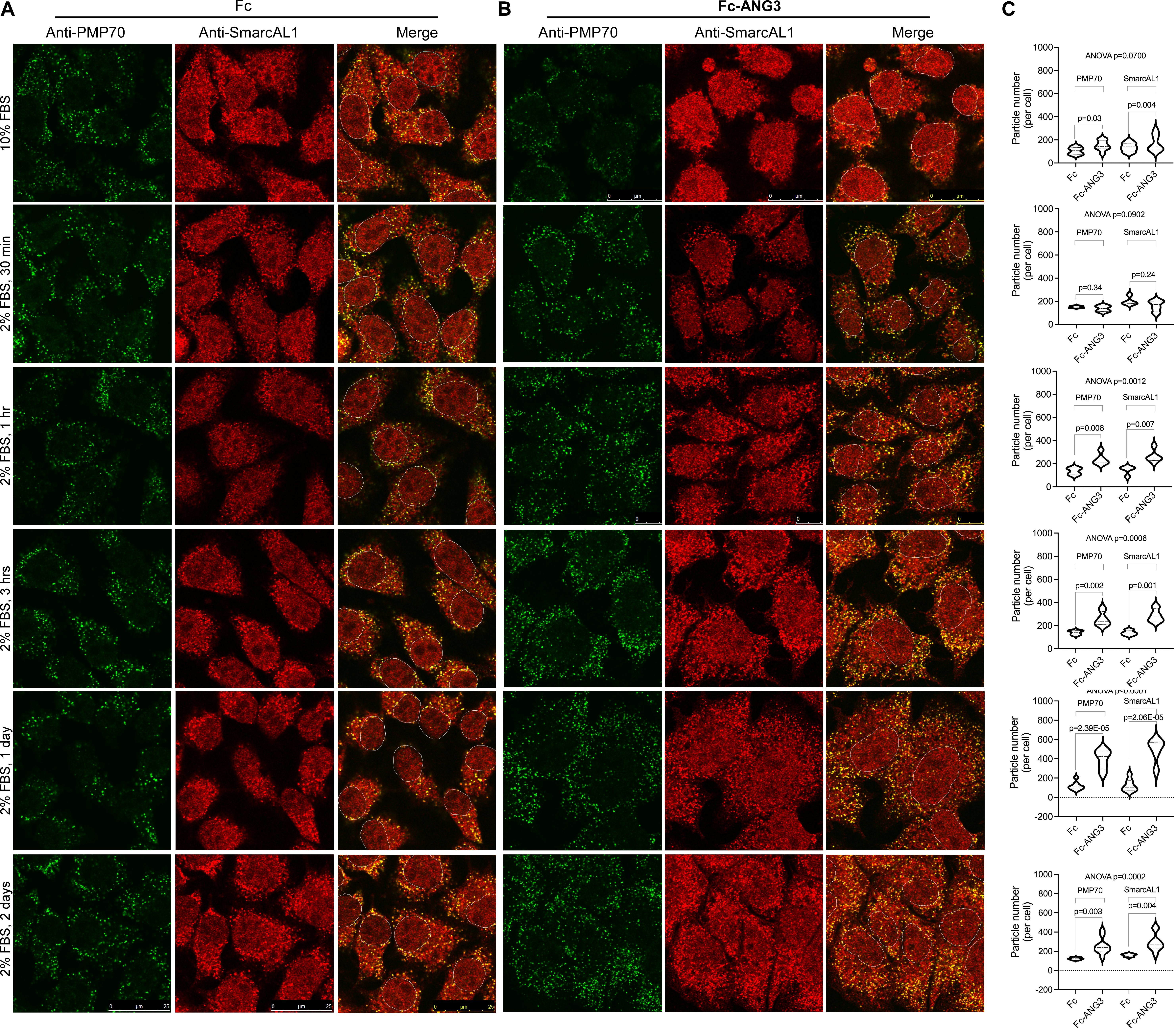

### Supplemental Fig. 6

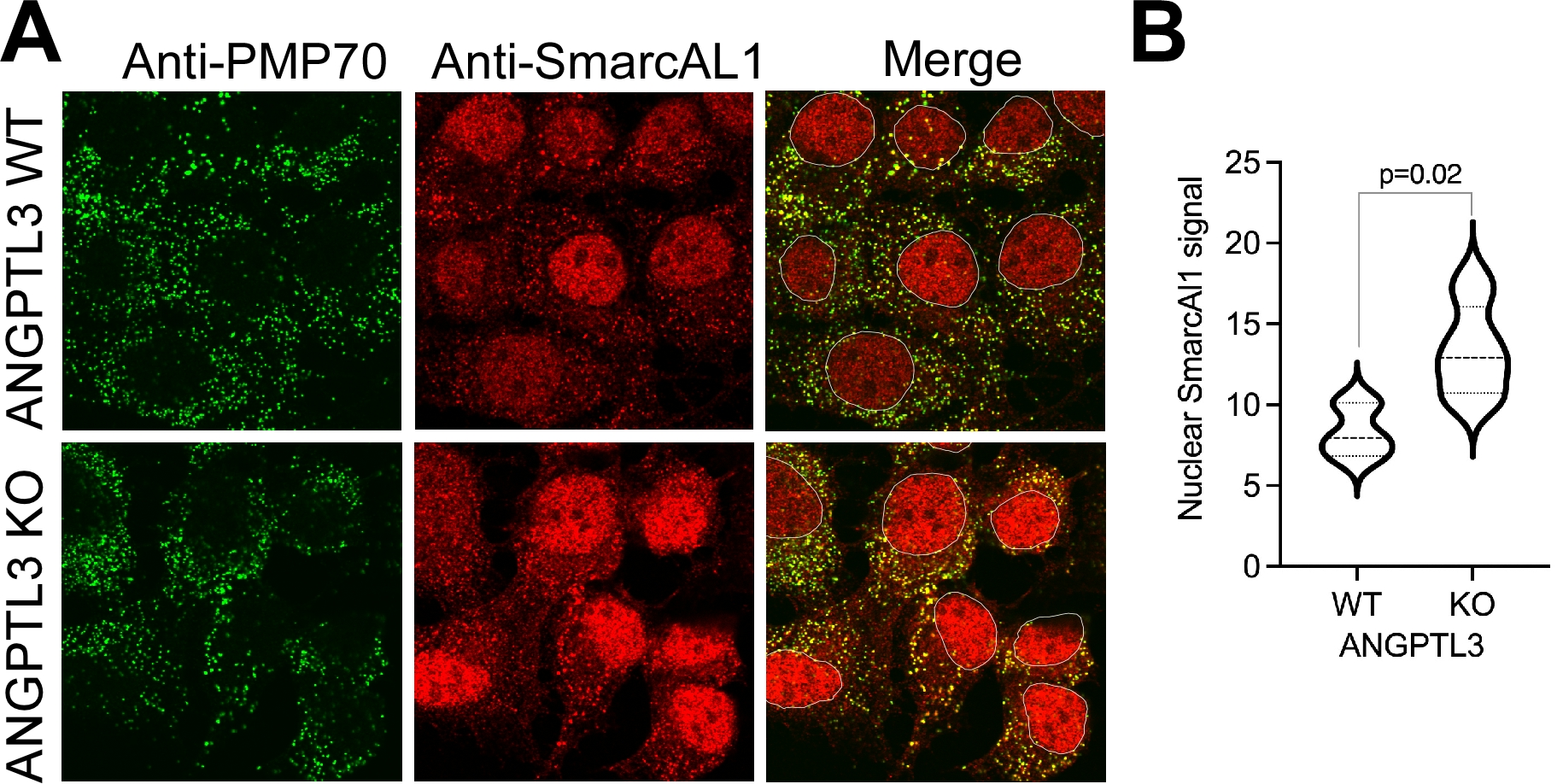

### Supplemental Fig. 7

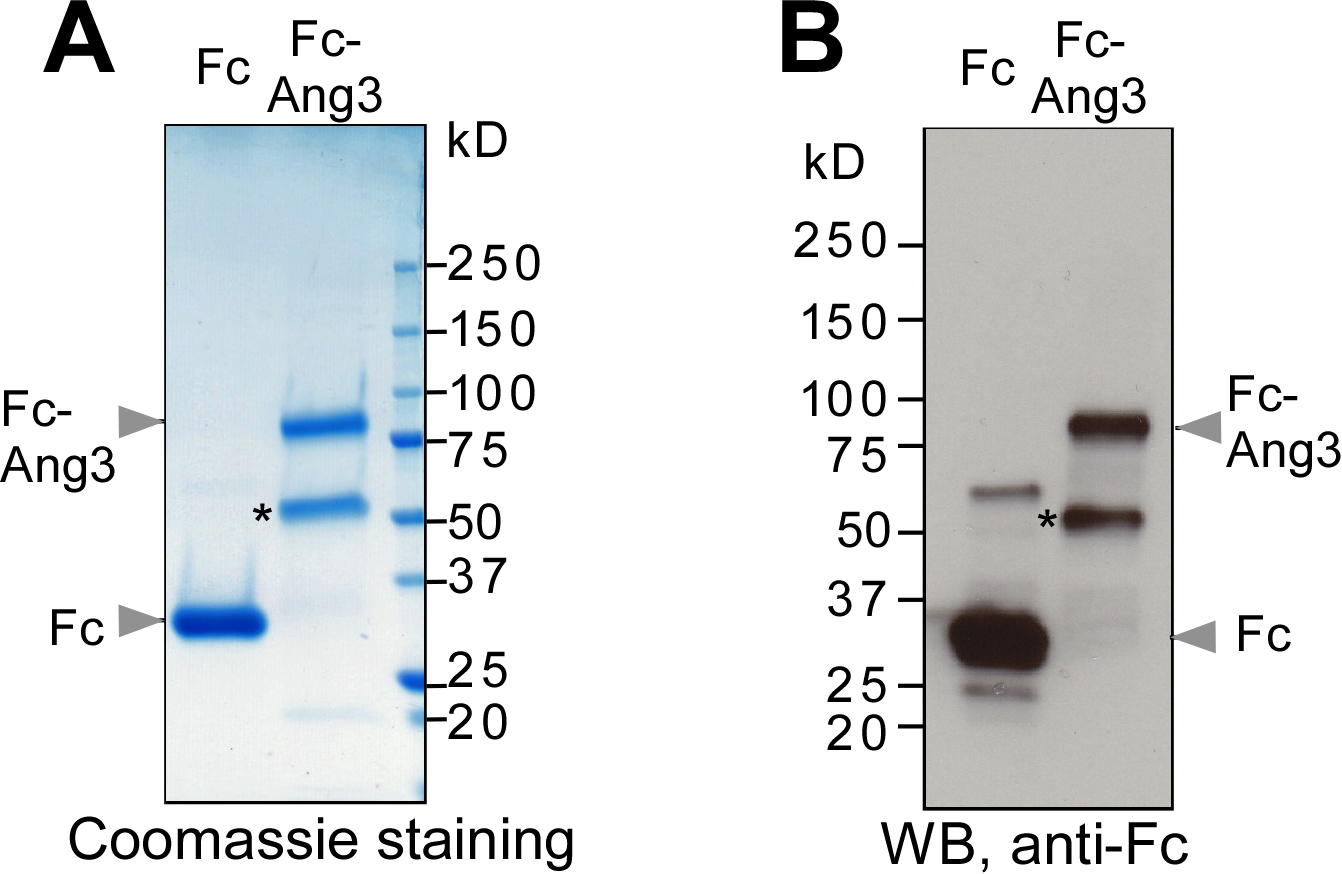
